# Nanoscale pattern extraction from relative positions of sparse 3D localisations

**DOI:** 10.1101/2020.02.13.947135

**Authors:** Alistair P. Curd, Joanna Leng, Ruth E. Hughes, Alexa J. Cleasby, Brendan Rogers, Chi Trinh, Michelle A. Baird, Yasuharu Takagi, Christian Tiede, Christian Sieben, Suliana Manley, Thomas Schlichthaerle, Ralf Jungmann, Jonas Ries, Hari Shroff, Michelle Peckham

## Abstract

We present a method for extracting high-resolution ordered features from localisation microscopy data by analysis of relative molecular positions in 2D or 3D. This approach allows pattern recognition at sub-1% protein detection efficiencies, in large and heterogeneous samples, and in 2D and 3D datasets. We used this method to infer ultrastructure of the nuclear pore, the cardiomyocyte Z-disk, DNA origami structures and the centriole.

## Introduction

Supramolecular complexes often form ordered structures, the organisation of which can yield insights into their cellular role. To reveal this organization, fluorescent labels can be targeted to specific molecules within a complex, then imaged with single molecule localisation microscopy (SMLM) approaches. SMLM is capable of achieving localisation precisions at or near the molecular scale, below 20 nm laterally and 50 nm axially^1^. When interpreting the resulting images, which are localisation maps, the intrinsic noise of SMLM arising from the stochastic switching of fluorophores can obscure the underlying molecular order. It is particularly difficult to uncover the underlying molecular organisation when a low fraction of the target molecules is localised with high precision, a common result of low labelling density or high background signal in 3D imaging^2^.

Single-particle averaging and reconstruction techniques have been developed and applied to enhance the signal-to-noise ratio and reveal underlying patterns of organisation from SMLM data^3–8^. Similarly, 1D and 2D autocorrelation (e.g. Fourier-domain processing) of SMLM reconstructions have been used to measure the periodicity in a biological structure^9, 10^. However, these methods require the target molecule to be efficiently labelled and detected with high signal to noise ratio, and the complex to be very highly ordered. To perform averaging, they also require either consistent orientation of the biological complex or classification and alignment of segmented regions of interest (ROIs), which presents further challenges^2^. Fourier analysis of SMLM data at high resolution requires the generation of ROIs with the same resolution (pixel size). Practically, this restricts the size and dimensionality of the input ROI (see **Suppl. Note 1**). These limitations restrict the applicability of existing methods to a small subset of supramolecular assemblies; thus, new approaches are required for pattern analysis and recognition of order in SMLM data.

## Results

We present a new fast and efficient method to assess any 2D or 3D SMLM dataset for potential regular structures through the analysis of relative positions of localisations. We name this PERPL analysis (pattern extraction from relative positions of localisations, **Fig. 1)**. This analysis markedly extends previous work using pair correlation^11–13^ by extending analysis into 3D and by comparing experimental inter-molecular distances against models of ordered and disordered macromolecular geometry, using appropriate statistical methods to help select the most probable model. This new method allows for rapid analysis of data, including large, 3D datasets in their entirety, at far greater resolution and with modest computational resources (see examples in **Suppl. Note1**) compared to Fourier analysis.

**Figure 1:**
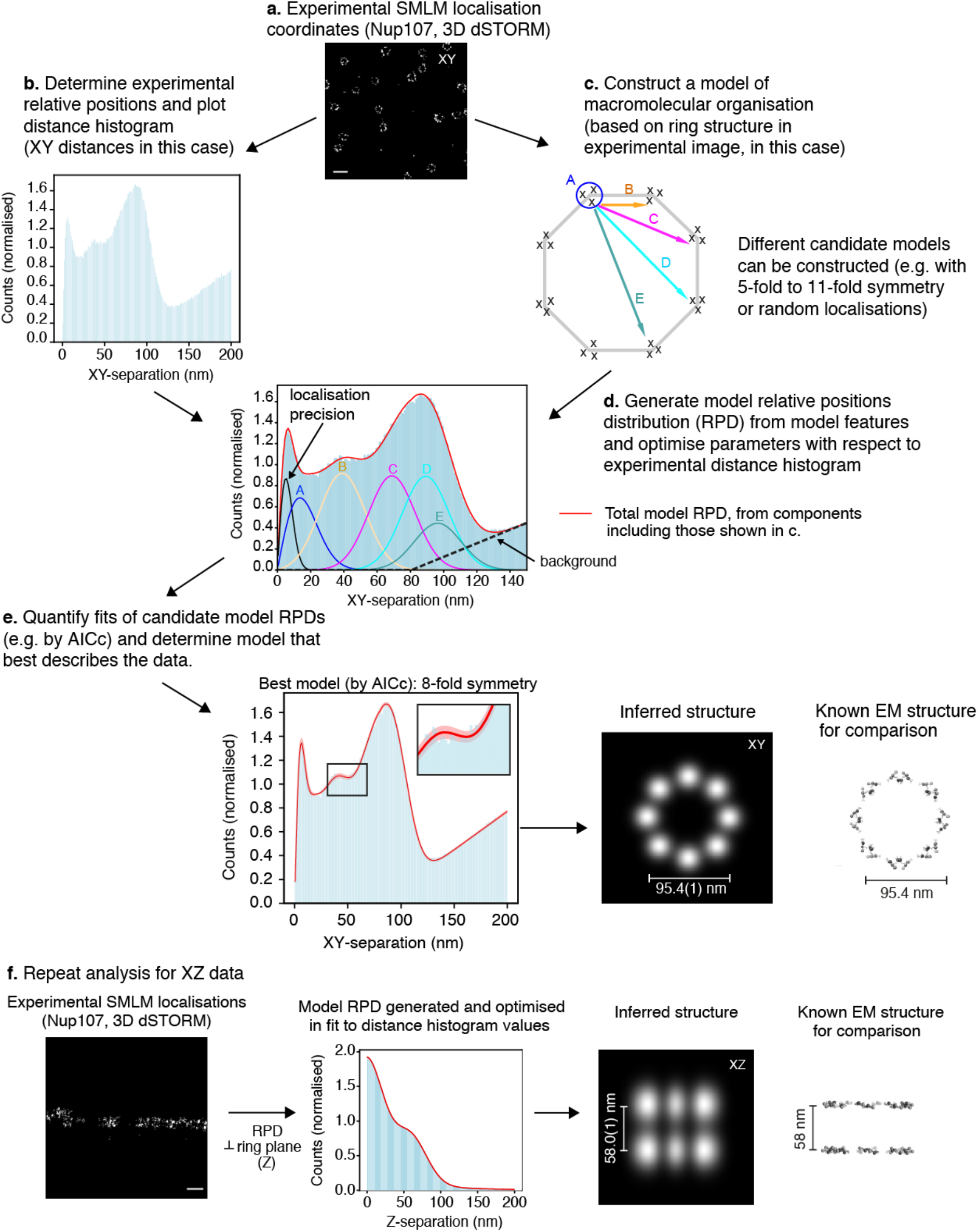
PERPL workflow and analysis of Nup107 localisations. Localisation coordinates from a 3D dSTORM experiment are rendered as a 2D image (**a**, scale bar: 200 nm), used to calculate a distance histogram (**b**, mean bin value scaled to 1.0), and to inform in silico candidate models of macromolecular geometry (**c**). The Nup107 XY candidate model contains inter-vertex distances (B-E), with vertices arranged symmetrically on a circle, components for repeated localisations of a single molecule (localisation precision) and unresolvable substructure in a cluster (A), and a background term (**c, d**). Discreet distances in the model (zero for localisation and clustering) are broadened using the distance distribution between two Gaussian clusters in 1, 2 or 3D^20^. The model relative position distribution (RPD) is the sum of these distributions (red line in **d**). The model RPD is fitted to the experimental distance histogram (**d**, **e**, pink is 95% confidence interval). The AICc values^19^ calculated for this and other models are compared to find the most likely model (**Suppl. Table 1**). The 8-fold model was the most likely for the Nup107 experimental data, in agreement with electron microscopy (EM) data^15^ (**e**). A two-layer model for Z-separations within the complex fitted well to the structure of the experimental distance histogram and also agreed with the EM data^15^ (**f**, scale bar: 200 nm, includes inferred structure in X in the XZ projection of the best model).

We validate and demonstrate this approach using a number of different datasets. First, we used a 3D SMLM dataset, containing images of nuclear pores in which Nup107 is labelled at the C-terminus^14^ (**Fig. 1a–f**). The arrangement of Nup107 is known from EM data^15^ providing us with a ground truth to validate our method. Nevertheless, we began our structural analysis using no prior information. The SMLM reconstruction shows many similar ring-like structures oriented with their axis of symmetry nearly aligned with the Z-direction (**Fig. 1a**). The first steps of the analysis are to take each localisation (*loc_ref*) across the entire FOV, select all localisations within a chosen distance of it (*locs_nearby*) and store the relative positions (RPs) in XYZ between each *loc_ref* and *locs_nearby*. We set a maximum pairwise distance of 200 nm, larger than the visible structures. This analysis uses the entire FOV and is fast, taking a few minutes to analyse a 16 × 17 × 0.9 μm volume containing 36k localisations on a standard laptop.

For the next step of the PERPL analysis, we calculated distance components across the XY plane from the 3D RPs and generated a histogram from them (**Fig. 1b**), called a relative position distribution (RPD). The appearance of peaks in this histogram implies an underlying set of characteristic separations, indicating that Nup107 is frequently found in specific geometric arrangements. To learn more about which architectures would be consistent with these separations, we generated in silico models for comparison. These candidate model structures were based on the observed organization (**Fig. 1a**) and thus had rotational symmetry. They were parameterised for diameter, degree of symmetry, localisation precision and a background distribution, and a substructure term to correct for molecules too close to resolve, or distributions of localisations resulting from overlapping images of nearby molecules^16^ (**Fig. 1c, d and Supplemental Fig. 1**). The RPD for each model was calculated and compared to that generated from the raw data (**Fig. 1d, e**), to obtain optimised model parameters.

To identify the model most likely to describe the real structure, or disorder, underlying the data we employed Akaike’s Information Criterion^17^ (AIC). Rather than categorically judging distributions as ‘significantly’ or ‘not significantly different’, AIC is a quantitative measure of information loss when approximating real data with a model (see **Online Methods**). The difference in AIC values between models also reflects the relative likelihoods (Akaike weights) that they capture the reality underlying the data^18^. We use the corrected AIC (AICc)^19^, which improves on the accuracy of AIC for more complex models evaluated at fewer data points.

In the example of Nup107, we tested models with rotational symmetries from 5- to 11-fold. The 8-fold symmetry model was selected as the most likely model, as it had the lowest AICc value and therefore the highest value for the Akaike weight (**Suppl. Table 1**). This model was approximately 4× more likely to correctly explain the distribution of the experimental data than the next most likely model (9-fold symmetry). The diameter of the 8-fold symmetric model was 95.4(1) nm (uncertainty is one s.d. on values of the fitted parameters; **Fig. 1e**, **Suppl. Table 2**), which fits well with that expected from EM data.

We then considered the axial structure of the complexes. Visual inspection of the XZ-reconstruction and the two lobes in the Z-distance histogram suggested a two-layer structure, where each layer has a spread in the Z-direction. A model of this kind, which included a Gaussian spread within each layer, resulted in a very good fit and indicated that Nup107 (C-terminus) would be found in layers separated by 58.0(1) nm (**Fig. 1f**, **Suppl. Table 3**), again agreeing well with EM data.

Following this result, we returned to our analysis of the XY-distances and now restricted it to pairs of localisations within only one of these layers, using axial distances ΔZ < 20 nm. Our revised analysis suggested that the 8-fold model is more than 10^10^× more likely to be the best model than the next closest (9-fold see **Suppl. Tables 4,5**), a much greater likelihood than the 4× obtained when all pairs of localisations with ΔZ < 200 nm were included. This indicates that each layer of the Nup107 complex is very likely to have an eight-fold symmetry in the XY plane, but that the simple model fits the data less well for the combination of the two layers. This may be explained by the known angular offset between the two layers^15^. Overall, PERPL analysis of the entire FOV of nuclear pores, naturally aligned in the XY plane, required no particle segmentation or registration and was in very good agreement with previous EM data^15^.

In our next example, we used PERPL to analyse 3D localisations of striated muscle Z-disk proteins (**Fig. 2a-f and Suppl. Figs 2-7**). Here, the choice of analysis dimensions is driven by lower-resolution features of the cardiomyocyte: its cylindrical symmetry, the cylindrical symmetry of its constituent myofibrils, and the highly regular repeating pattern of Z-disks (∼100nm wide, ∼2 μm apart), which span the width of the cell^21, 22^. Once again, prior knowledge of the molecular-scale structure was not used for this analysis. However, the tetragonal lattice arrangement of actin filaments and α-actinin-2 (ACTN2) is known and presents a challenging validation structure with characteristic distances under 20 nm^21, 22^ (**Fig. 2d,f**). Both limited labelling density and high background in such a thick, dense structure result in a low fraction of localisations per target molecule.

**Figure 2:**
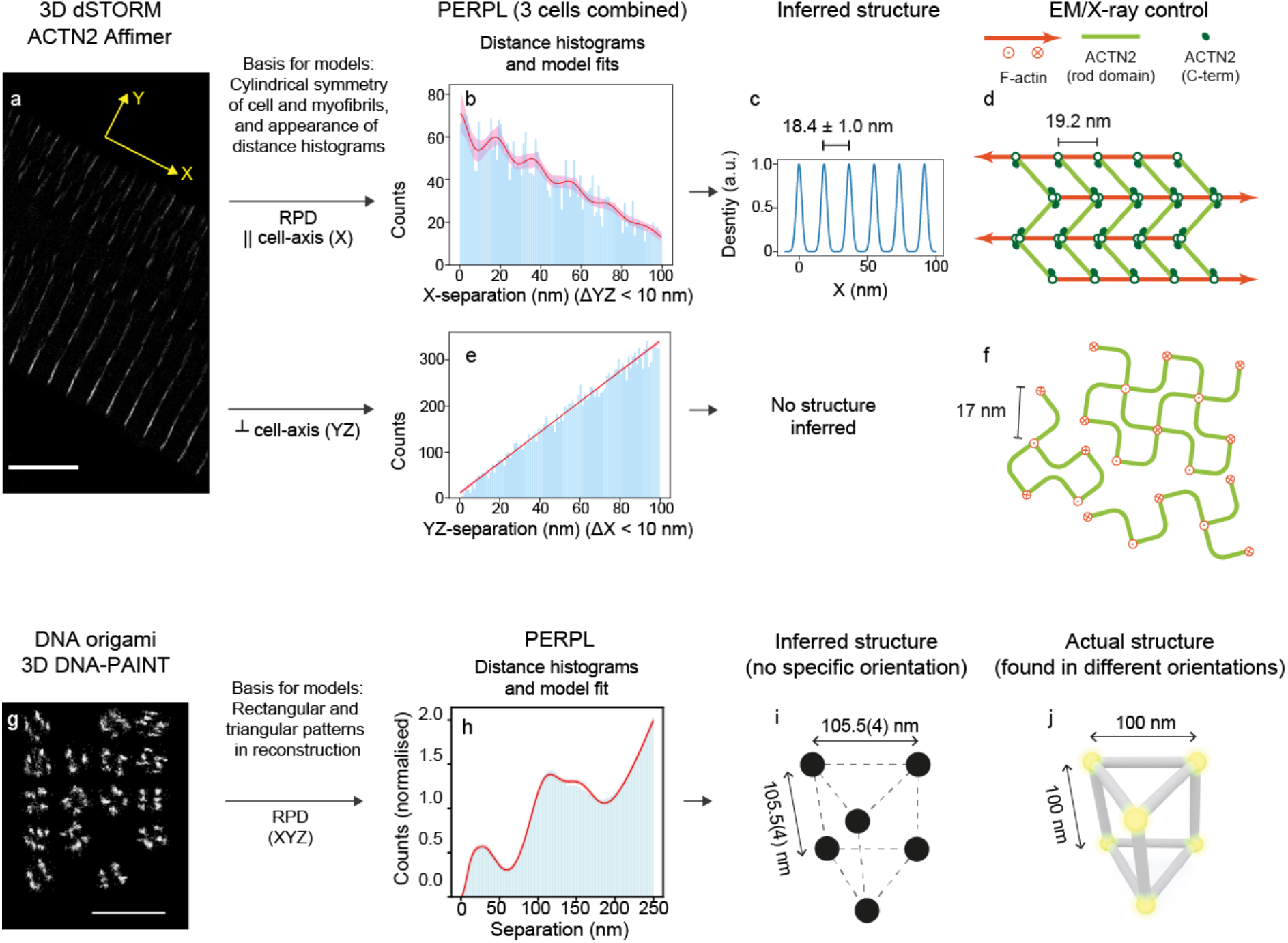
PERPL analysis of 3D ACTN2 and DNA-origami localisations. In analysis of 3D dSTORM localisations of ACTN2 (**a**, scale bar: 5μm), the histogram of separations in X supported a model RPD resulting from a linear repeating structure (**b**, **c**). The inferred model matched previous results from other modalities (**d**)^21^. The best-supported model to the distance histogram in YZ was an isotropic, homogeneous distribution (**e**). No ordered geometry was inferred, and we did not detect the known square lattice features in the plane of the Z-disk^21, 23^, which is known to contains domains in different orientations (**f**)^21^. We analysed a list of 3D DNA-PAINT^24^ localisations of a DNA-origami sample (**g**, scale bar: 500 nm), without knowledge of its structure. The appearance of a reconstruction of the FOV (**g**) suggested models of simple geometries on a square lattice. We used the Euclidean distance between localisations in 3D for the model RPD (**h**), to allow the models to be correct for all possible orientations of the structure. The best supported model was a triangular prism (**h**, **i**), which was the correct structure (**j**), and on which we inferred close to the correct side-length (**i**, **j**).

We analysed the distribution of experimental 3D dSTORM localisations (see **Methods**) of an Affimer ^**25**^ raised against the CH domains (calponin homology domains: the C-terminal actin-binding site^26^) of ACTN2 and directly conjugated to Alexa Fluor 647 (see **Online Methods, and Suppl. Fig. 3**). X-ray crystallography of the Affimer confirmed its binding to isolated CH2 domains (Suppl Fig. 3, Suppl. Table 6). Relative positions and distance histograms over three cells were aggregated together to perform this analysis. First, we addressed the question of organisation in the axial direction of the cells (**Fig. 2a-d**). The smoothed distance histogram (**Suppl. Fig. 4**) suggested that the structure may contain a repeating distance. To test this, we constructed models for localisations found at multiples of a unit distance along the cell-axis (**Suppl. Fig. 5**), together with a model for a purely random distribution within the thickness of the Z-disk and one containing repeated localisations of the same molecule. AIC analysis demonstrated that the best model of the localisation distribution included an instance of the ACTN2-CH2 domain every 18.4(1.0) nm along the cell-axis with 97% confidence (**Suppl. Table 7 and legend, Suppl. Table 8**). This repeating distance compared well with the 19.2 nm periodicity of ACTN2 binding sites obtained from EM^22^.

Our current understanding of the Z-disk in cardiac muscle suggests that it is composed of four to six layers of ACTN2 crosslinks^21^. Using AICc analysis to compare Z-disk models with four, five or six layers (**Suppl. Table 7**), showed that a model describing five layers (**Suppl. Table 8**) had the highest relative likelihood, although the differences between these models were not great enough to robustly select one model over the others. This may be due to natural variability of the Z-disk, but the analysis may also be limited by the quality of the data. Specifically, at greater distances across the finite (∼100 nm) Z-disk, the number of RPs obtainable, and therefore the signal to noise ratio, is reduced. In the transverse (YZ) plane, the relative imprecision of localisations in Z, and the misalignment of Z-disk domains **(Fig. 2)**, prevented the robust observation of characteristic features.

We also performed PERPL analysis on 3D PALM data of fluorescent protein (FP) fusions to three Z-disk proteins: mEos2:ACTN2, Lim and SH3 domain protein-2:mEos3.2 (LASP2:mEos3.2), and myopalladin:mEos3.2 (MYPN:mEos3.2) (**Suppl. Fig. 6**). The arrangements of LASP2 (N-terminus) and MYPN (N-terminus) in the Z-disk are as yet poorly understood, although their C-termini are both known to bind at ACTN2 (C-terminus)^27, 28^.

After filtering for localisation precision (< 5 nm), these datasets were limited to < 1% protein detection efficiencies, leading to lower signal to noise in the distance histograms (**Suppl. Fig. 7**). In this situation a kernel density estimate (KDE) of the RPD can be used. The KDE of the ACTN2:mEos2 axial data was fitted well by the same repeating density model, with a repeat of 20.20(8) nm, similar to that observed for the 3D dSTORM Affimer data. The KDE of the LASP2 data contains peaks that also appear to be separated by a similar distance. However, the peaks in the optimised RPD model for both the LASP2 and MYPN data were too broad to observe such a repeat, and the optimised model parameters had very high uncertainties (**Suppl. Fig. 7, Suppl. Table 9**). Therefore, while these models did result in lower AICc values than a linear fit, inspection of the model RPDs, parameter values and uncertainties does not allow us to be confident about any repeating structure that might be present.

As a third example, we carried out blind analysis on localisations of a DNA-origami structure (**Fig. 2g-j**). Reconstruction of the FOV revealed geometric structures in different orientations on an approximately square grid. Therefore, we constructed in silico models of simple geometric structures and included features reflecting the presence of localisations at nearby grid points. Use of the total distances between localisations in XYZ led us to conclude that the most probable model to explain the data is a triangular prism structure with sides of equal length on a square grid (**Fig 2g–j, Suppl. Fig. 8**). This model matches the design of the DNA origami^29^; furthermore, we obtained a side-length estimate of 105.5(4) nm, compared to the design length of 100 nm. The triangular prism was convincingly the most likely model (**see Suppl. Tables 10,11**), although there were deviations between the data and the model, and the estimate of the side-length was slightly high. We attribute these discrepancies partly to the assumption of isotropic localisation precision used in the model, which while not strictly true for the data, allowed us to conveniently average over all orientations. Second, the proximity of the structures at adjacent square lattice points is likely to result in an extra distribution of characteristic distances that were not accounted for, and which would require a more complex model.

In these analyses, we have presented examples where we reduce the dimensionality of the SMLM data (e.g. from 3D coordinates to a pair of 1D distance histograms), in order to account for more orientations of a complex in a model RPD (**Figs. 1,2**). However, it is important to note that the dimensionality of the original data is retained as the RPs are calculated between 3D localisations. The PERPL analysis can also be developed to use 3D and 2D distance histograms (**Suppl. Fig. 9**) for more complex visualisation and analysis.

Finally, we tested the use of PERPL analysis to define the relative quality of particles to be selected for other analysis methods. To do this, we used SMLM data on centriolar protein Cep152, in which the particles (centrioles) had already been segmented from an SMLM image and classified according to their orientation^8^. We analysed each ‘top-view’ particle, and scored for uncertainty in fitted model parameters associated with distances between clusters in a 9- fold symmetric model. This allowed us to discard ill-defined particles before averaging them together, a process which improved the quality of particle averages (**Suppl. Fig. 10**).

## Discussion

PERPL analysis therefore contributes to the development of tools needed to explore and assess patterns in SMLM data. It allows quantitative, accurate assessments of 2D and 3D model structures without requiring particle segmentation and alignment, including when the fraction of molecules localised is very low (< 1%), and with only modest computational resources. It can also be used to assess the quality of particles segmented from an SMLM dataset. In most, but not all, of these examples, the analysis has taken advantage of commonalities in orientation between instances of a complex, which can occur naturally or be selectively obtained by segmentation and classification. However, as SMLM techniques continue to improve 3D localisation precision and particle segmentation, we anticipate that further developments of PERPL analysis will also enable more detailed quantitative assessment of 3D model structures, against sparse localisations within randomly oriented biological complexes. Finally, the application of PERPL need not be restricted only to SMLM data. It can be used to analyse organisation of any data where localisation coordinates are obtained from image features, such as clusters, or nodes in a network structure.

To enable the community to conveniently use this analysis method on other protein arrangements, we have provided the Python scripts used here as **Supplementary Software** and at https://bitbucket.org/apcurd/curd_et_al_perpl_software/src/master/. Examples of output distance histograms and fits generated by the software are shown in **Suppl. Fig. 11,12**. Test data is also provided at https://bitbucket.org/apcurd/curd_et_al_perpl_test_data/src/master/.

## Acknowledgements

The work was funded by the UK Medical Research and the UK Biotechnology and Biological Sciences Research Councils (MR/K015613/1 and BB/S015787/1 (to MP and A.Curd) and BB/M011151/1 DTP studentship to BR); the Wellcome Trust (Institutional Strategic Support Fund at University of Leeds to RH and A.Cleasby and travel award to MP); the Intramural Research Program of the National Heart, Lung and Blood Institute; and the National Institute of Biomedical Imaging and Bioengineering, US National Institutes of Health. The dSTORM system was funded by alumnus M. Beverley, in support of the University of Leeds ‘making a world of difference’ campaign. R.J. acknowledges support by the DFG through the Emmy Noether Program (DFG JU 2957/1-1), the ERC through an ERC Starting Grant (MolMap, Grant agreement number 680241), the Max Planck Society, the Max Planck Foundation and the Center for Nanoscience (CeNS). T.S. acknowledges support from the DFG through the Graduate School of Quantitative Biosciences Munich (QBM). Isolated cardiomyocytes were a kind gift from the Steele Group, University of Leeds. We thank Ulf Matti and Philipp Hoess for sample preparation and imaging of the Nup107 cells, and Niccolò Banterle for the centriole sample preparation. We would also like to acknowledge Michael W. Davidson for his contributions to the development of the mEos3.2 constructs.

## Author contributions

A.Curd conceived and implemented the analysis approach, and developed software. J.L. developed software. R.H. and A.Cleasby imaged Z-disk proteins in 3D PALM. B.R. imaged ACTN2 Affimer in 3D dSTORM. C.H and B.R. crystallised the Affimer-CH domain construct and solved its structure. M.B. and M.P. developed fluorescent protein constructs. H.T., H.S. and M.P. developed 3D PALM labelling and imaging techniques. C.S and S.M. provided the Cep152 localisation data and conceived the particle quality assessment concept. J.R. provided the 3D Nup107 localisation data. M.P. conceived the Z-disk protein experiment and acquired confocal data on Z-disk protein labelling. A.Curd and M.P. wrote the manuscript with input from all other authors.

## Methods

### 3D dSTORM localisations of SNAP-tagged Nup107

Data was acquired as described in Li et al.^1^. In short, U2OS cells that expressed Nup107–SNAP^2^ were fixed and labelled with benzylguanine-Alexa Fluor 647 (NEB, Ipswich, MA, USA) and imaged on a custom microscope in a standard blinking buffer (50 mM Tris, pH 8, 10 mM NaCl, 10% (w/v) d-glucose, 35 mM 2-mercaptoethylamine, 500 μg/mL glucose oxidase, 40 μg/mL catalase, 2 mM cyclooctatetraene. Single molecules were localized using a Gaussian PSF model and the data were drift corrected using redundant cross-correlation.

We filtered the localisations for high precision, less than 10 nm uncertainty in Z according to the MLE fitting routine^1^.

### Affimer to ACTN2

To raise Affimers to ACTN2, the CH domains from human ACTN2 were expressed and purified from E.Coli as described^3^, biotinylated, and used in a phage-display screen as described^4^. All 7 unique Affimers obtained from the screen were expressed and purified, and directly dye labelled (either using Alexa Fluor 647, or Alexa Fluor 488) using the unique C-terminal cysteine as described^5^. The dye-labelled Affimers were then tested for their ability to label Z-discs in fixed adult rat cardiomyocytes by confocal microscopy. Those that did were then tested in dSTORM (using Alexa Fluor 647 labelled Affimers). The Affimer that demonstrated the best labelling for dSTORM was then used in subsequent experiments.

### Protein crystallization, data collection and structure determination

Crystals were obtained at 20°C by the sitting-drop vapour diffusion method using 20% (w/v) polyethylene glycol 3350 and 0.2 M ammonium tartrate with a protein concentration of 8 mg ml^−1^. The crystals were flash-cooled in liquid nitrogen after soaking for 30 seconds in mother liquor solution containing 25% (v/v) glycerol as a cryo-protectant. X-ray diffraction data were collected at the Diamond Light Source on beamline I04 to 1.2 Å resolution at 100 K. The diffraction images were indexed and integrated using DIALS^6^ before subsequent scaling in AIMLESS^7^ and data processing in the CCP4i2 suite^8^.

The unit cell parameters for the crystal are a=46.2Å, b=48.5Å, c=147.0Å, α=β=γ=90.0° in space group ***P***2_1_2_1_2_1_ with one ACTN2:AF9 complex in the asymmetric unit cell. The structure was determined by molecular replacement using the program PHASER^9^ with the human Calponin domain of Alpha Actinin-2 structure (PDB code 5A38^3^) and the truncated Affimer (PDB code 4N6T^10^) as the search models. Initial rounds of automated model building were performed in BUCCANEER^11^ followed by iterative rounds of manual model building in COOT^12^ and refinement using REFMAC5^13^. During the course of the model building structural validations were carried out using the program MOLPROBITY^14^. The N-terminal 16 residues and C-terminal 13 residues of the human Calponin domain were disordered and were not included in the final refined structure. The structure factor and coordinate files have been deposited in the Protein Data Bank with accession code 6SWT.

### mEos Z-disk adenovirus constructs

Three Z-disk protein constructs were made. The α-actinin-2 mEos2 construct was the same as previously described^3^, in which mEos2 was fused to the C-terminus of α-actinin-2 (mEos2:ACTN2(C)). LASP2(N):mEos3.2 (LASP2 is LIM and Src homology 3 Protein-2, also known as LIM-nebulette) was generated from a GFP-LASP2 construct kindly provided by Carol Gregorio and as originally reported by Terasaki et al.^15^. To generate LASP2(N):mEos3.2, the LASP2 cDNA was subcloned into pdc315 (Microbix) in frame with the N-terminal Eos3.2 coding sequence. Similarly, MYPN(N):mEos3.2 was generated from a GFP-myopalladin construct kindly provided by Siegfried Labeit^16^. Adenovirus was generated from each of the constructs as previously described^3, 17^, purified (Vivapure AdenoPACK 100, Sartorius) and titred using a TCID_50_ assay, before aliquotting and storage at −80°C.

### Isolated cardiomyocyte culture

Coverslips were coated with 50 μg/ml laminin overnight at 37°C. For 3D PALM, we used 25-mm diameter, #1.5 (Scientific Laboratory Supplies, MIC3350) coverslips, cleaned as by Shroff et al.^18^. Laminin was removed and coverslips allowed to dry. Cardiomyocytes were collected in Tyrodes buffer, media was exchanged for cardiomyocyte media (M199 media, Gibco, supplemented with 5 mM creatine, 5 mM taurine, 2 mM Na pyruvate, 2 mM L-carnitine, 0.1 μM insulin, 250 μg normocin, 40 μM cytochalasin D) and cells were seeded onto laminin coated coverslips and allowed to attach for 2 hrs at 37°C. For 3D dSTORM, the cells were immediately fixed and stored at 4°C prior to staining. For 3D PALM, media was changed and the cells infected with adenovirus constructs in fresh medium at MOIs of x100. The cardiomyocytes were then incubated for 24–40 hrs at 37°C before fixing.

### Immunostaining cardiomyocytes for confocal imaging

Isolated rat cardiomyocytes for confocal imaging were plated onto 13-mm diameter washed glass coverslips in medium. mEos-tagged Z-disk constructs were introduced via adenoviral expression, before fixation with 4% PFA for 20 min. Cells were permeabilised with 0.1% Triton X-100 in PBS for 10 min, washed gently with PBS and blocked with 3% bovine serum albumin (BSA) for 1 hr. Fixed cardiomyocytes were co-stained with anti-ACTN2 antibody (1:400) (Sigma, A7811, lot no. 024M4758), anti-titin antibody T12^19^ (Boehringer Mannheim, discontinued), which recognises Z-disk titin epitopes, followed by staining with Alexa Fluor 647-labelled donkey anti-mouse secondary antibody (1:100, ThermoFisher). All antibodies and dyes were diluted in 0.2% BSA. Coverslips were washed and mounted in Prolong Gold Antifade. Cells were imaged using a 63x objective (NA 1.4), on a Zeiss 880 LSM Airyscan confocal microscope.

### Fixation and processing of cardiomyocytes for 3D PALM or dSTORM

Cells were fixed with final concentration of 2% paraformaldehyde (PFA) in phosphate buffered saline (PBS) for 10 min. Fixed cells were washed three times for 5 min with PBS prior to 3D PALM. Cells were imaged within 24-48 hrs. For dSTORM, after fixation, the cells were permeabilised with 0.5% Triton X 100 in PBS for 5 minutes, blocked with 5% bovine serum albumin (BSA) in PBS for 1 hour and then stained using the ACTN2 Affimer labelled with Alexa-647, stock Affimer was diluted 1/750 in PBS with 1% BSA to give a final concentration of 0.6 μg/ml, for 60 mins at room temperature. Cells were then washed in PBS and stored briefly at 4°C before 3D dSTORM imaging.

### 3D PALM Instrumentation for Z-disk imaging

3D PALM^20^ used an inverted microscope (Olympus, IX81) with a 60x, 1.2 NA, water-immersion objective lens (Olympus, UPLSAPO60XW). An automated *x-y* stage with additional piezoelectric adjustment in *z* (PZ-2000, Applied Scientific Instrumentation) was fitted to accommodate a focus lock (C-focus, Mad City Labs) and the sample. The coverslips were mounted in chambers (I-3033-25D, Applied Scientific Instrumentation), held in a stage insert (I-3033, Applied Scientific Instrumentation). Lasers at 561 nm and 405 nm (Jive, Cobalt and LuxX, Omicron, respectively, integrated in a LightHUB, Omicron) provided widefield excitation and photo-activation of mEos constructs, together with a custom-built 2x beam expander before the rear illumination port of the microscope. A multi-band excitation filter (zet405/488/561/640m, Chroma) was used with a multi-band beamsplitter (zt405/488/561/640rpc, Chroma) and a multi-band emission filter (zet488/561/640m-TRF). The imaging path included a 1.6x magnifier internal to the microscope, an external 1.2x magnifier (Diagnostic Instruments, DD12BXC) and a cylindrical lens with *f* = 150 nm (Thorlabs, LJ1629RM-A), which provided an astigmatic point-spread function for a single emitter^21^. Images were captured by a back-illuminated, electron-multiplying CCD camera, cooled to −80°C (Andor Technology, iXON Ultra, model DU-897U-CSO-#BV), using published scripts^20^ called from the camera interface (Andor Technology, SOLIS).

### Z-disk protein localisation with 3D PALM and dSTORM

The data acquisition workflow^20^ included capture of calibration images of a gold nanoparticle (742031, Sigma-Aldrich) in steps of 50 nm in *z* over a 4 μm range and use of these particles for drift-tracking. The mEos2 and mEos3.2 FP tags were stochastically photoswitched using 405-nm illumination and excited with 561-nm illumination to provide sparsely distributed emission events from the ensemble of the labels. The Alexa Fluor-647 labelled Affimer to ACTN2 was imaged in imaging buffer composed of PBS, pH 8.0 with 10% glucose (w/v), 0.5 mg/ml glucose oxidase, 80 μg/ml catalase, 110mM β-mercaptoethanol and the fluorophore was excited using a 642-nm laser (100 mW), with EMCCD gain 150, and the blinking rate controlled by gradually increasing 405 nm laser power from 2-20 mW. At least 11,000 frames were collected at 20 Hz and acquisition was stopped when the number of emission events per frame became negligible. Jump tracking was used to image the fiducials at the coverslip surface as well as blinks within the cell at a different height to fit within the calibration series.

Emission events were localised using *palm3d*^20^, with those lasting for more than one frame averaged to provide one ‘linked’ localisation. Localisations were then binned into a histogram for display, accounting for drift and *x-y* distortion by the cylindrical lens. In order to make the localisation data available for analysis, new Python scripts were required, which are available at https://bitbucket.org/apcurd/palm3d_extra/. These scripts provide users of *palm3d* with corrected localisation data, including a precision estimate, allowing filtering and subsequent analysis of the dataset.

We used the maximal area of the field of view that did not include localisations of the gold nanoparticles, as there are many repeated localisations of these. When initially included in the analysis, distances between these repeated localisations overwhelmed the relative position distribution in later analysis.

Localisation precision was estimated as the standard error on the localisation position (σ / √N), using a 2D Gaussian fit of the relevant image from the *palm3d* calibration stack (σ, mean of two widths from the 2D fit) and the photon count above background (N, available from the *palm3d* data for every linked localisation). This simple precision estimate is likely to be an under-estimate^22^, but was useful as a method to allow filtering for higher-precision localisations and to assist in the resolution of short distances within the complex (e.g. ∼20-nm distances between ACTN2 localisations). 1.3 × 10^5^ localisations over 6 cells were included in the mEos2:ACTN2 analysis, after filtering for estimated localisation precision within 5 nm.

### 3D DNA-PAINT localisations of DNA-origami structures

#### DNA origami structure assembly

DNA origami nanostructures were assembled in a one-pot reaction in a final volume of 50 μl. The assembly mix contained p8064 single-stranded DNA scaffold strand (tilibit nanosystems) at a final concentration of 10 nM, single-stranded core DNA oligonucleotides at 100 nM, DNA-PAINT P1 (5’-Staple-TT ATACATCTA-3’) docking sites at 500 nM, biotinylated DNA strands at 800 nM in a buffer of 5 mM Tris and 1 mM EDTA supplemented with 12 mM MgCl_2_. The pooled strand solution was heated to 80 °C for 5 min followed by a thermal ramp from 60 °C to 4 °C over the course of 17 h. Assembled nanostructures were purified by agarose gel electrophoresis (1.5 % (w/v) agarose, 0.5×TAE, 10 mM MgCl_2_, 1×SYBR Gold) at 3 V/cm for 3 h at 4 °C. Gel bands were cut, crushed and structures were purified with Freeze ‘N Squeeze spin columns (Bio-Rad) for 5 min at 1,000×g at 4 °C.

#### Sample preparation and DNA-PAINT imaging

Flow chambers for imaging were assembled with a coverslip (no. 1.5, 18×18 mm^2^) attached to a standard microscopy glass slide with two strips (approximately ∼0.5 cm – 1 cm apart) of double-sided sticky tape. As a first step, 20 μl of biotin-labeled bovine serum albumin (1 mg/ml, dissolved in buffer A (10 mM Tris-HCL, pH 7.5, 100 mM NaCl and 0.05% (v/v) Tween 20, pH 7.5)) was flown into the flow chamber and incubated for 2 min. Afterwards, the chamber was washed with 40 μl of buffer A, followed by incubation with 20 μl streptavidin (0.5 mg/ml in buffer A) and incubated for another 2 min. After washing with 40 μl of buffer A and 40 μl of buffer B (5 mM Tris-HCl, pH 8, 10 mM MgCl_2_, 1 mM EDTA, 0.05 % (v/v) Tween 20 at pH 8). 200 pM of the self-assembled DNA origami structure was incubated in the channel and allowed to attach for 2 min. The chamber was washed with 40 μl of buffer B and finally 3 nM Cy3b labeled DNA-PAINT imager strand (CTAGATGTAT-Cy3b) was added in buffer B supplemented with a PCA/PCD/Trolox oxygen scavenging system. The chamber was sealed with picodent before imaging. Nanostructures were imaged for 15000 frames, with an exposure time of 200 ms per frame and at a laser excitation (560 nm) intensity of 3 kW/cm^2^.

#### Microscopy Setup for DNA-PAINT

DNA-PAINT imaging was carried out on an inverted Nikon Eclipse Ti microscope (Nikon Instruments) with the Perfect Focus System, applying an objective-type TIRF configuration with an oil-immersion objective (Apo SR TIRF 100×, NA 1.49, Oil). A 561 nm excitaion laser (200 mW, Coherent Sapphire) was used. The laser beam was passed through a cleanup filter (ZET561/10x, Chroma Technology) and coupled into the microscope objective using a beam splitter (ZT561rdc, Chroma Technology). Fluorescence light was spectrally filtered with an emission filter (ET600/50 and ET575lp for 561 nm excitation, Chroma Technology) and imaged on a sCMOS camera (Andor Zyla 4.2) without further magnification, resulting in an effective pixel size of 130 nm after 2×2 binning. A cylindrical lens was inserted into the beampath in front of the camera for 3D imaging, the corresponding calibration was performed as previously reported^21^. Camera Readout Sensitivity was set to 16-bit, Readout Bandwidth to 200 MHz.

#### DNA-PAINT image reconstruction

DNA-PAINT images were reconstructed with the Picasso software suite as previously reported^23^. Single structures were picked and aligned in their native orientation in a two-dimensional grid for further processing. This grid of structures was the FOV analysed with PERPL (**Fig. 2g–j**), without prior knowledge of the origami structure or its arrangement. Localisations were not filtered for estimated localisation precision in this case.

### 2D dSTORM localisations of Cep152

Human centrioles were purified and antibody labelled as described previously^24^. STORM imaging of immunostained centriole samples was performed using a recently developed flat-field epi illumination microscope^25^. Briefly, a 642 nm laser (2RU-VFL-P-2000-642-B1R, MPB Communications) was used to switch off fluorophores on the sample, while a 405 nm laser (OBIS, Coherent) controlled the return rate of the fluorophores to the fluorescence-emitting state. A custom dichroic (ZT405/561/642/750/850rpc, Chroma) reflected the laser light and transmitted fluorescence emission before and after passing through the objective (CFI60 PlanApo Lambda Å∼60/NA 1.4, Nikon). After passing the emission filter (ET700/75M, Chroma), emitted light from the sample was imaged onto the sCMOS camera (Prime, Photometrics). Axial sample position was controlled using the pgFocus open hardware autofocus module (http://big.umassmed.edu/wiki/index.php/PgFocus). Typically, 40,000 frames at 10 ms exposure time were recorded using Micromanager^26^. Imaging was performed using an optimized STORM buffer as described previously^27^. Image stacks were analysed using a custom CMOS-adapted analysis routine^28^. Localisations from individual centrioles were segmented and extracted using SPARTAN as described previously^24^. We filtered the localisations for high estimated localisation precision, within 5 nm.

### Relative positions of localisations

We obtained relative positions within a distance filter chosen to be larger than the relevant features observed in images. Among the Nup107 localisations, we obtained all relative positions in the FOV within 200 nm in X, Y and Z. Within the cardiomyocyte Z-disk protein data, we cropped each FOV to exclude the repeated localisations of gold nanoparticles. For all remaining localisations in each FOV, we obtained all relative positions within 200 nm in X, Y and Z. In the DNA-origami data, we obtained all relative positions in the FOV within 250 nm in X, Y and Z. In the Cep152 data, we obtained all relative positions within 1000 nm in X and Y in each pre-segmented particle.

### Representation of inferred models of Nup107 organisation and electron microscopy map (Fig. 1e, f)

We plotted the points according the inferred model (**Fig. 1**, **Suppl. Table 2**) and applied Gaussian smoothing according to the fitted broadening parameters in the models. The electron microscopy map was generated in the UCSF Chimera package from the Computer Graphics Laboratory, University of California, San Francisco (supported by NIH P41 RR-01081)^29^.

### Aggregation of relative positions of Z-disk proteins from multiple cardiomyocytes

3D SMLM cardiomyocyte reconstructions, and the corresponding sets of 3D relative localisation positions, were aligned by first rotating an XY-projection of each cell reconstruction (e.g. **Fig. 2a**) such that the cell-axis pointed along X. The corresponding set of 3D relative localisation positions was then rotated by the same angle. Thus, 3D relative position data was in the same coordinate system across all cells for aggregation. Relative position data were aggregated from 3 cells for the ACTN2 Affimer analysis, from 6 cells for mEos2:ACTN2, 5 cells for LASP2:mEos3.2 and 5 cells for MYPN:mEos3.2.

### Z-disk protein distance histograms

The X distance histogram (**Fig. 2b**) contains the X component of the 3D relative positions with a component in the YZ plane < 10 nm. The YZ distance histogram (**Fig. 2e**) included relative positions with a component in X < 10 nm. The 2D (X vs. YZ) distance histogram (**Supplementary Fig. 9c**) was normalised in the cell-transverse direction according to the radial distribution function for a 2D homogeneous, isotropic localisation distribution.

For the 2D and 3D distance histograms (**Suppl. Fig. 9**), relative position data from the Z-disk proteins was smoothed with a Gaussian kernel with σ = 4.8 nm, reflecting the uncertainty on measuring the distance between two points which themselves have an uncertainty of 3.4 nm (the mean XY precision estimate of the filtered localisations included in the mEos2:ACTN2 analysis).

### Construction of candidate models for macromolecular complex geometry

Typically, the coordinates of proteins in a candidate model for a complex were calculated in a function wherein the distances between them would be determined by input variables. These variables would later be used by the fitting function to vary the geometry generating the relative position distribution (RPD) that was fitted to the experimental distance histograms. Example of such models are found in 2D in the function *generate_polygon_points()* in *modelling_general.py*, and in 3D in *polyhedramodelling.py*.

### Generation of candidate model relative position distributions (RPDs) for macromolecular complexes

Model RPDs were typically generated by calculating the positions of localisations in hypothetical models of macromolecular complex geometry, as in the function *rot_sym_only* in *rot_2d_symm_fit.py*. The 1D, 2D or 3D distance distribution between clusters with a Gaussian distribution^30^ was applied to the relevant distances in a candidate model to give a realistic, parametric distribution of relative positions. Other components of models included a spread representing repeated localisations of the same molecule and a spread representing unresolvable substructure or a distribution of mislocalisations resulting from overlapping images of emitters. Examples of these components can also be found in *rot_sym_with_replocs_and_substructure_isotropic_bg* in *rot_2d_symm_fit.py.* We also used model RPDs for background in SMLM data, examples of which can be found in *background_models.py*, and the functions *rot_sym_with_replocs_and_substructure_isotropic_bg* and *rot_sym_replocs_substructure_isotropic_bg_with_onset* in *rot_2d_symm_fit.py*. The latter is the RPD model used in the XY analysis of **Fig. 1** and accompanying text and Supplementary Information. Noise in the localisation data (e.g. due to finite localisation precision or residual inaccuracy in frame-to-frame drift correction, after drift correction has been applied to the SMLM image sequence) in general increases the broadening parameters in the model. Where localisation precision is estimated in a model, this includes the effect of such noise.

Starting values and bounds for parameters (e.g. distances, broadening on the distribution) are also required by the model-fitting implementation. Example of these, including default values and opportunities to edit them, are available, for instance, for the rotationally symmetric models of *rot_2d_symm_fit.py*, in the functions *create_default_fitting_params_dicts, set_up_model_replocs_substruct_iso_bg_with_onset_with_fit_settings*, and *set_up_model_replocs_substruct_no_bg_with_fit_settings*.

### Distance histogram fitting and comparison of models

Experimental distances between localisations in the chosen directions were binned into a histogram, with which model relative position distributions (RPDs) could be compared. In cases of low SNR (**Suppl. Fig. 7**), we used the kernel density estimate of the relevant distances as described. The model RPDs were fitted to the experimental distance distribution in Python scripts with *scipy.optimize.curve_fit*^31^, a least squares fitting function. This function outputs the covariance matrix for the optimised model parameters. Uncertainties on parameters (1 s.d.) are √(diagonal elements of the covariance matrix). Confidence intervals on the model RPD values at each distance are calculated from the covariance matrix and derivatives with respect to the model variables, which are calculated with *numdifftools*, a package based on the MATLAB toolbox DERIVESTSuite (John D’Errico, 2006, http://www.mathworks.com/matlabcentral/fileexchange/13490-adaptive-robust-numerical-differentiation). This calculation can be found in the function *stdev_of_model* in *modelling_general.py*. Bounds on the 95% confidence intervals are the estimated model RPD value ± 1.96 × s.d. calculated in this way.

We used Akaike’s Information Criterion (AIC)^32, 33^ to calculate differences in the level of support that the experimental data give to the candidate model RPDs (relative likelihoods of the models, given the experimental data). A lower AIC value for a model means that less information is lost when approximating the data with that model. The AIC includes a penalty for increasing the number of parameters (*K*) in a model, and so does not automatically favour complex models that over-fit the data. Specifically, we used a corrected AIC (AICc) that adjusts the penalty for increasing *K*, as *K* becomes non-negligible with respect to the number of data points (e.g. distance histogram bins)^33, 34^. Differences in AIC (AICc) values are used in calculations of relative likelihood that the candidate model RPDs are correct, given the data^28, 30^: likelihood ∝ exp {−1/2 AICc}^33, 35^; likelihood ratios resulting from differences in AICc values are given as ‘Akaike weights’, in a sum to 1, to aid interpretation^33^.

## Data availability

The datasets generated and analysed in this study are available from the authors upon reasonable request.

## Code availability

The code developed in this study is available at https://bitbucket.org/apcurd/curd_et_al_perpl_software/src/master/, and test data for the software can be found at https://bitbucket.org/apcurd/curd_et_al_perpl_test_data/src/master/. Updated versions of the code can be found at https://bitbucket.org/apcurd/perpl-python3/.

