## supplemental figures and data for "Nanoscale pattern extraction from relative positions of sparse 3D localisations"

**Supplementary Note 1: Comparison of PERPL analysis to Fourier-analysis of SMLM reconstructions.**

We chose a small region of interest (ROI) with which to compare the resources required for analysis with fast Fourier transforms (FFT) and PERPL. The DNA-PAINT localisations of **Figure 2** span a volume with dimensions (2.301 × 1.023 × 0.796 $\mu m^3$). Generation via FFTs of the autocorrelation that would contain comparable distance information to the histogram of **Figure 2** would require an initial 3D reconstruction (3D histogram) of the localisation data with 1-nm voxel size, corresponding to 2301 × 1024 × 796 voxels. 64-bit information is used in standard implementations of FFT calculations, including the Python numpy library[1], a freely available library with which we developed the PERPL analysis software. Construction of this 3D array, requiring ~15.0 GB of memory, was not possible with the computing resources used for all of the reported PERPL analysis (8 GB RAM, quad-core CPU processing at 2.10 GHz). Calculation of the FFT would double this memory requirement by adding an imaginary component to each pixel value in the 3D array. Use of the FFT to calculate the autocorrelation subsequently requires the concurrent use of at least one additional 3D array with complex values (calculations based on the complex conjugate and inverse FFT), increasing the memory requirement to ~60.0 GB. This estimate assumes no padding, that would further increase the memory requirement, is required to prevent artifacts resulting from the assumption that the input array is periodic beyond its boundaries.

Construction of the 3D histogram was possible with a 2-nm voxel size, but calculation of the autocorrelation was not, with the same computing resources mentioned above. Construction of the 3D histogram and calculation of the 3D autocorrelation was possible with a 5-nm voxel size. Generation of the total XYZ distance histogram as used in this analysis would require further work on summing over spherical shells in the autocorrelation output.

Non-equispaced FFT (NFFT) implementations exist[2], which may use sparse data as input, and may be suitable for SMLM localization coordinates. However, these are more complex than the FFT to understand and apply in autocorrelation analysis. The memory and number of arithmetic operations required for NFFT scale in a similar way to the FFT, with the spatial frequency bandwidth required, or equivalently the resolution of spatial information desired in a final autocorrelation output[2]. Subsequent calculation of the autocorrelation would then require the same amount of available memory as with FFT.

On the other hand, with PERPL analysis, we were able to compute $1.0 \times 10^7$ 3D pairwise relative positions (RPs) within 250 nm in XYZ, using all 14,219 localisations in the dataset, in 38 minutes. We use the filter distance (250 nm in this case) to dramatically reduce memory requirements compared with computation and storage of all RPs in a FOV. No precision limit is imposed up to this point; relative positions retain the precision stored by the SMLM localization technique, without binning into reconstruction voxels as required for the FFT. Bin size for the distance histograms can then be chosen to be arbitrarily small, depending on the requirements of the analysis. It is also possible to use smaller portions of the localization data (e.g. using one out of every two localisations) to speed up the calculations, or if a memory limit is reached. This decreases signal-to-noise ratio in the resulting distance histograms, but characteristic distances underlying the data remain the same, and models can be fitted as usual to the resulting distance histograms. These histograms from reduced data can still contain information from millions of relative positions.

The Nup107 data of **Figure 1** covered a much larger ROI, at 16.4 × 17.4 × 0.870 µm. This is ~130× the volume of the DNA-PAINT ROI, requiring the corresponding increase in memory for FFT calculations. With PERPL analysis, we analysed all localisations in the dataset with estimated localization precision $\sigma_Z < 10$ nm, to reduce broadening and improve resolution of any underlying structure in the distance histograms. We computed $1.0 \times 10^6$ 3D pairwise relative positions within 200 nm in XYZ, using all 36,297 localisations, in 11 minutes.

For many samples, PERPL analysis is therefore more practical than high-resolution FFT analysis.

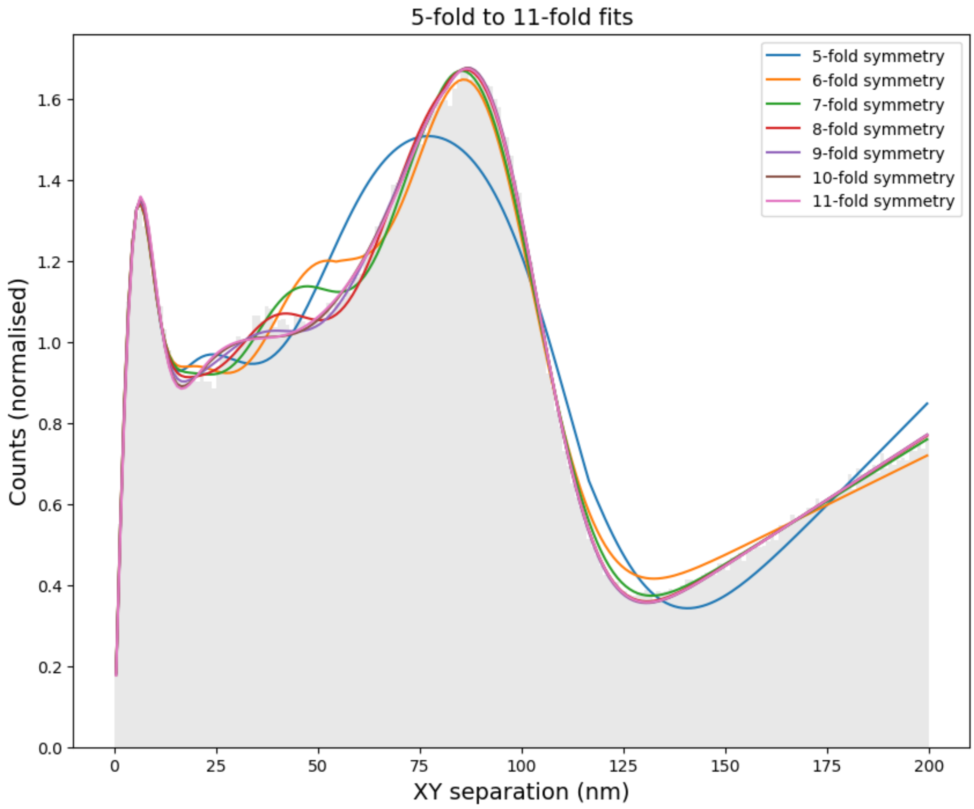


**Supplementary Figure 1: Best fits of candidate relative position distributions (RPDs) to the experimental RPD of SNAP-tagged Nup107 in Fig. 1, for different degrees of rotational symmetry.** Quantitative fit results are found in Suppl. Table 1. This plot was generated and saved by *rot_2d_symm_fit.py* in Supplementary Software (see also **Suppl. Fig. 12**).

**Supplementary Table 1: Corrected Akaike Information Criterion (AICc) values[3] and relative likelihoods (Akaike weight, summing to one)[4] that the RPD models are the closest to the true molecular RPD, for pairwise XY-distances between Nup107 localisations (Fig. 1 and Suppl. Fig. 1).** The eight-fold rotational symmetric model has approximately 4× stronger support than the next most likely model (nine-fold rotational symmetry).

| Rotational symmetry order | AICc | Akaike weight |
|---|---|---|
| 5 | -911.79 | $1.3\times10^{-137}$ |
| 6 | -1168.39 | $6.8\times10^{-82}$ |
| 7 | -1337.76 | $4.1\times10^{-45}$ |
| 8 | -1541.69 | 0.79‡ |
| 9 | -1539.08 | 0.21 |
| 10 | -1484.65 | $3.2\times10^{-13}$ |
| 11 | -1469.28 | $1.5\times10^{-16}$ |

‡ Selected model.

**Supplementary Table 2: Parameters and uncertainties (1 s.d.) for the best fit RPD model (8-fold) to pairwise distances between Nup107 localisations.** The background term (see **Fig. 1d**) is an approximation reflecting the fact that the ring-like image features (nuclear pores), may be near to each other but do not overlap. The background is expected to have a linear form (isotropic in 2D) beyond the diameter of a nuclear pore, dominated by the distribution of other nuclear pores. It must also have a lower rate of increase at smaller distances, which we have approximated to zero, up to an 'onset' distance.

| | | |
|---|---|---|
| **Localisation precision term** | Amplitude | 7.4(2) a.u. |
| | Spread (precision, s.d.) | 3.63(5) nm |
| **A (cluster term)** | Amplitude | 15.3(2) a.u. |
| | Spread (s.d. of approx. Gaussian cluster) | 9.6(1) nm |
| **B–E (rotational symmetry contributions)** | Diameter of structure | 95.4(1) nm |
| | Amplitude of contributions | 15.73(5) a.u. |
| | Spread (broadening of contributions, s.d.) | 14.18(7) nm |
| **Background term** | Onset | 80(1) nm |
| | Gradient | 0.00645(9) a.u. $nm^{-1}$ |

**Supplementary Table 3: Parameters and uncertainties for the two-layer RPD model for pairwise distances in Z between Nup107 localisations.** The model contains two layers of localisations, each with the same Gaussian spread, and an exponentially decaying background. This background distribution for ΔZ has the same form as the distribution in Z of the probability of excitation in the evanescent illumination field of the TIRF instrumentation[5].

| | | |
|---|---|---|
| **Two-layer components** | Separation between layers | 58.0(1) nm |
| | Spread of layer (s.d.) | 15.92(4) nm |
| | Within-layer term: amplitude | 46.8(8) a.u. |
| | Between-layer term: amplitude | 39.6(5) a.u. |
| **Background term** | Amplitude | 0.21(3) a.u. |
| | Exponential scale parameter | 71(5) nm |

**Supplementary Table 4: Akaike information and weightings (summing to one) of the relative likelihood of RPD models to be the closest to the true RPD, for pairwise XY-distances between Nup107 localisations assumed to be within the same layer of the Nup107 complex (ΔZ < 20 nm).** The eight-fold rotational symmetric model is far more strongly supported than any other model. The difference between this and the closer results for all pairwise XY-distances (Suppl. Table 1, not filtered by ΔZ) can be understood to be due to imperfect registration of the two layers of the Nup107 complex, smearing out the overall XY-RPD.

| Rotational symmetry order | AICc | Akaike weight |
|---|---|---|
| 5 | -862.91 | $9.6\times10^{-99}$ |
| 6 | -1074.15 | $7.1\times10^{-53}$ |
| 7 | -1219.24 | $2.3\times10^{-21}$ |
| 8 | -1314.29 | 1.0‡ |
| 9 | -1250.73 | $1.6\times10^{-14}$ |
| 10 | -1206.15 | $3.3\times10^{-24}$ |
| 11 | -1198.05 | $5.7\times10^{-26}$ |

‡ Selected model.

**Supplementary Table 5: Parameters and uncertainties for the best fit RPD model (8-fold) to pairwise distances between Nup107 localisations assumed to be within the same layer of the Nup107 complex (ΔZ < 20 nm).**

| | | |
|---|---|---|
| **Localisation precision term** | Amplitude | 18.8(4) a.u. |
| | Spread (precision, s.d.) | 3.64(3) nm |
| **A (cluster term)** | Amplitude | 16.2(4) a.u. |
| | Spread (s.d. of approx. Gaussian cluster) | 9.1(2) nm |
| **B–E (rotational symmetry contributions)** | Diameter of structure | 95.5(1) nm |
| | Amplitude of contributions | 15.52(8) a.u. |
| | Spread (broadening of contributions, s.d.) | 13.5(1) nm |
| **Background term** | Onset | 83(2) nm |
| | Gradient | 0.0056(2) a.u. $nm^{-1}$ |

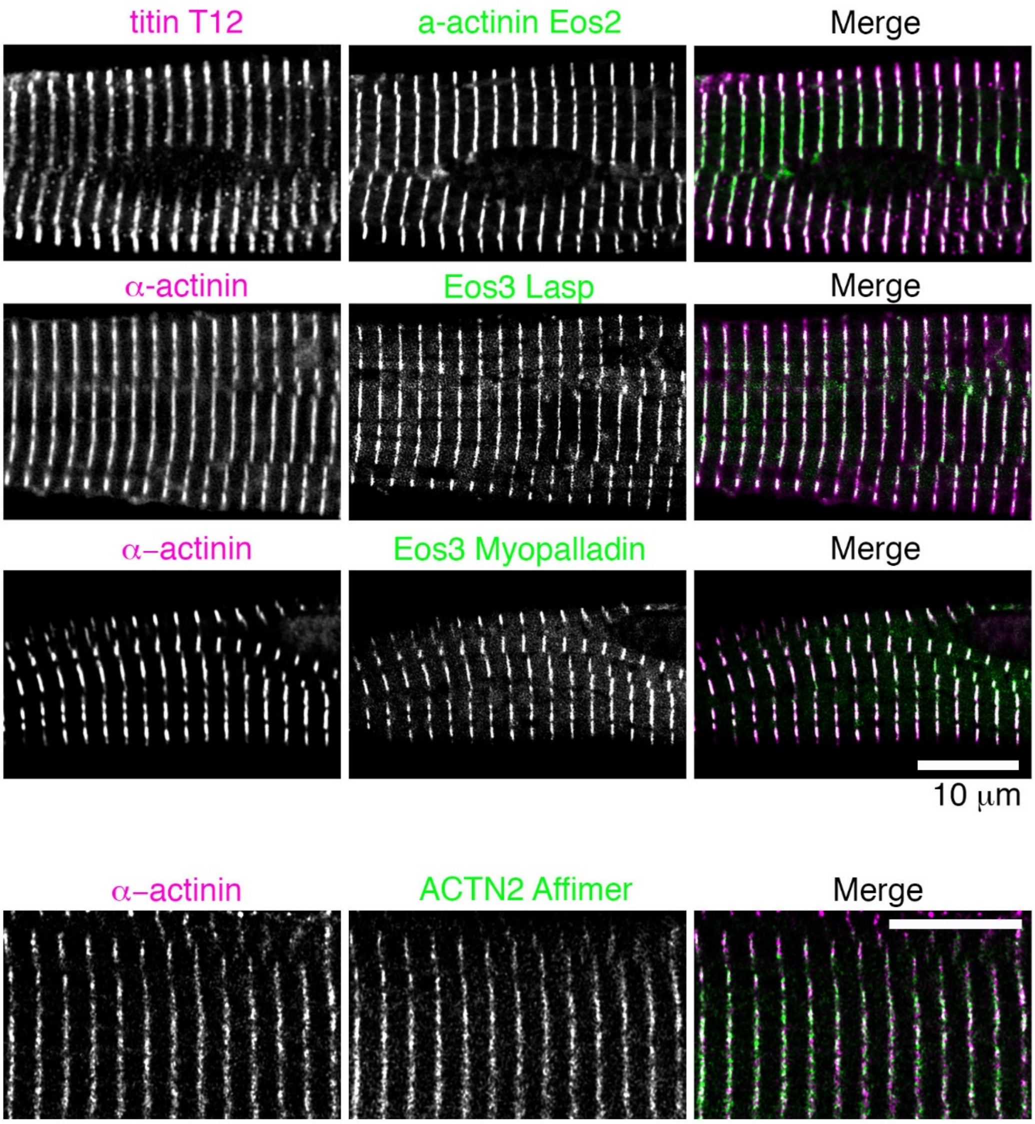


**Supplementary Fig. 2: Fluorescent protein (FP) constructs and ACTN2 Affimers localise to the Z-disks of cardiomyocytes.** Isolated cardiomyocytes expressing the FP constructs were fixed, co-stained for Z-disk markers, and imaged in confocal microscopy (antibodies specific to ACTN2 or Z-disk titin (T12)), or isolated cardiomyocytes (not expressing FP constructs) were fixed and stained with the ACTN2 Affimer and the Z-disc marker for α-actinin, and imaged using confocal microscopy. The Z-discs can be seen spanning the width of the cell. Scale bar is 10 μm for all images.

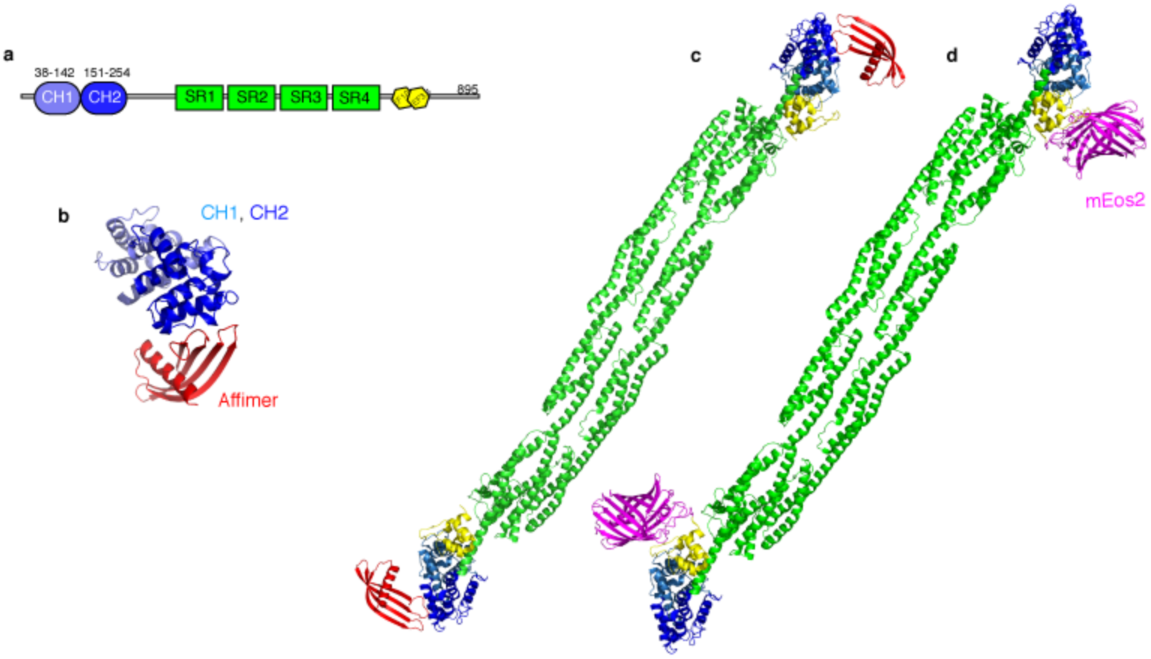


**Supplementary Fig. 3: Structure of the Affimer bound to the CH domains (PDB 6SWT).** (**a**) Schematic for α-actinin-2, showing the different domains – CH1, CH2 (which bind to F-actin), the spectrin repeats 1-4 (SR1, SR2, SR3 and SR4) followed by the EF hands. (**b**) Affimer (red) bound to the second CH domain, in the crystal structure (PDB 6SWT). The Affimer is ~2nm in size. (**c**) The Affimer in context – bound to the α-actinin-2 dimer. (**d**) For comparison, the position of mEos2, which is fused to the C-terminus of α-actinin-2, is shown.

**Supplementary Table 6.** Data collection, processing and refinement statistics for α-actinin-2:Affimer complexes.

| | ACTN2:AF9 |
|---|---|
| Source | Diamond beamline i04-1 |
| Wavelength (Å) | 0.9159 |
| Resolution range (Å) | 73.50–1.20 (1.23–1.20) |
| Space group | $P2_12_12_1$ |
| Unit-cell parameters (Å) | *a*=46.2, *b*=48.5, *c*=147.0 |
| Completeness (%) | 98.9 (93.3) |
| No. of observed reflections | 603488 (23921) |
| No. of unique reflections | 103011 (7078) |
| Redundancy | 5.9 (3.4) |
| $< I/\sigma(I) >$ | 10.4 (1.6) |
| Wilson B factor | 13.5 |
| $R_{merge}$ (%)§ | 7.3 (80.2) |
| $R_{pim}$ (%)¥ | 2.9 (49.3) |
| $CC_{1/2}$ | 0.99 (0.62) |
| Refinement statistics | |
| Resolution range for refinement (Å) | 73.5–1.2 |
| *R* factor (%) | 14.9 |
| $R_{free}$ (%)† | 16.4 |
| No. of protein non-H atoms | 1848 |
| No. of water molecules | 230 |
| R.m.s.d bond lengths (Å) | 0.008 |
| R.m.s.d bond angles (˚) | 1.4 |
| Average overall *B* factor ($Å^2$) | |
| All atoms | 22.9 |
| Protein | 22.1 |
| Water | 32.0 |
| Ramachandran analysis, the percentage of residues in the regions of plot (%) ‡ | |
| Favoured region | 97.5 |
| Outliers | 0.3 |
| PDB code | 6SWT |

*Values given in parentheses correspond to those in the outermost shell of the resolution range.

§ $R_{merge} = \sum_{hkl}\sum_i \left| I_i(hkl) - \langle I(hkl)\rangle \right| / \sum_{hkl}\sum I_i(hkl)$

¥ $R_{pim} = \sum_{hkl}\{1/[N(hkl)-1]\}^{1/2}\sum_i \left| I_i(hkl) - \langle I(hkl)\rangle \right| / \sum_{hkl}\sum_i I_i(hkl)$

† $R_{free}$ was calculated with 5% of the reflections set aside randomly.

‡ Ramachandran analysis using the program MolProbity [6].

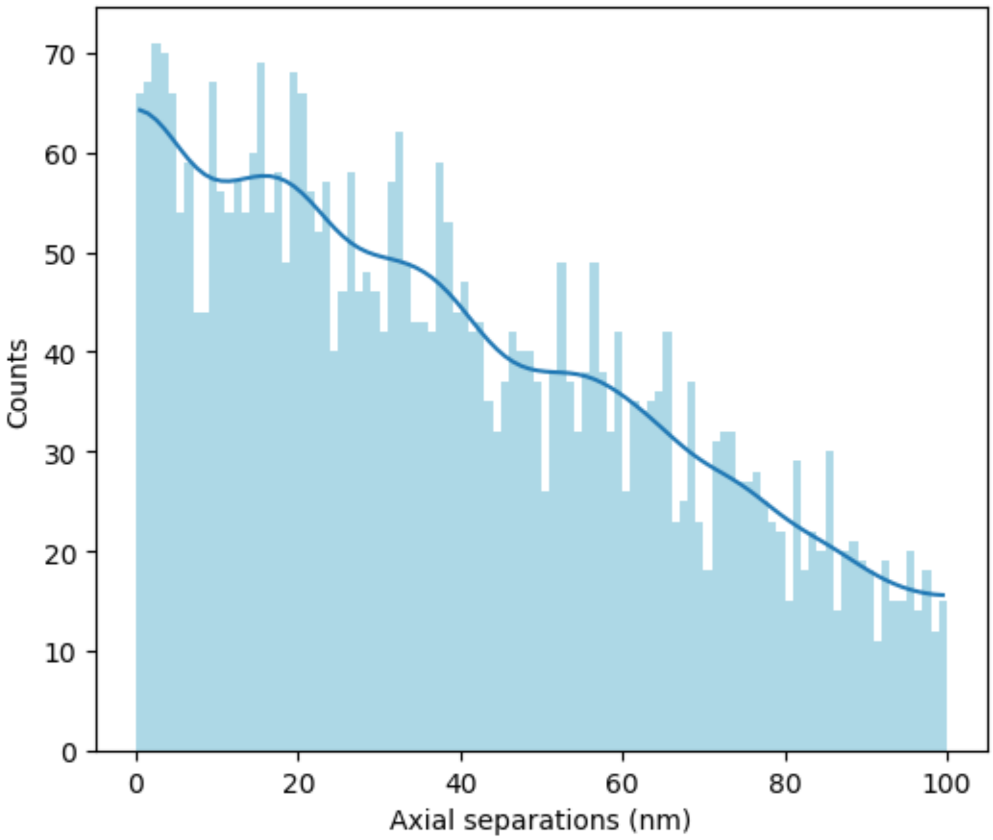


**Supplementary Fig. 4: Histogram and smoothed profile for distances along the cell-axis between ACTN2 Affimer localisations in rat cardiomyocytes, aggregated over 3 cells.** Gaussian filter with $\sigma = 4.4$ nm applied to the bin values ($\sqrt{2} \times$ average localisation precision estimate for included localisations, see Online Methods). There is a general descending linear trend, expected for localisations confined within the thickness of the Z-disk (~100 nm). The smoothed profile also hints at peaks occurring at a repeating distance, just under 20 nm. A feasible biological structure can be proposed (or in this case is already known from EM), that would explain such peaks, in which ACTN2 binds at a regular repeating distance along the actin filaments. The likelihood of that model, relative to the linear trend alone, can be assessed using the AICc.

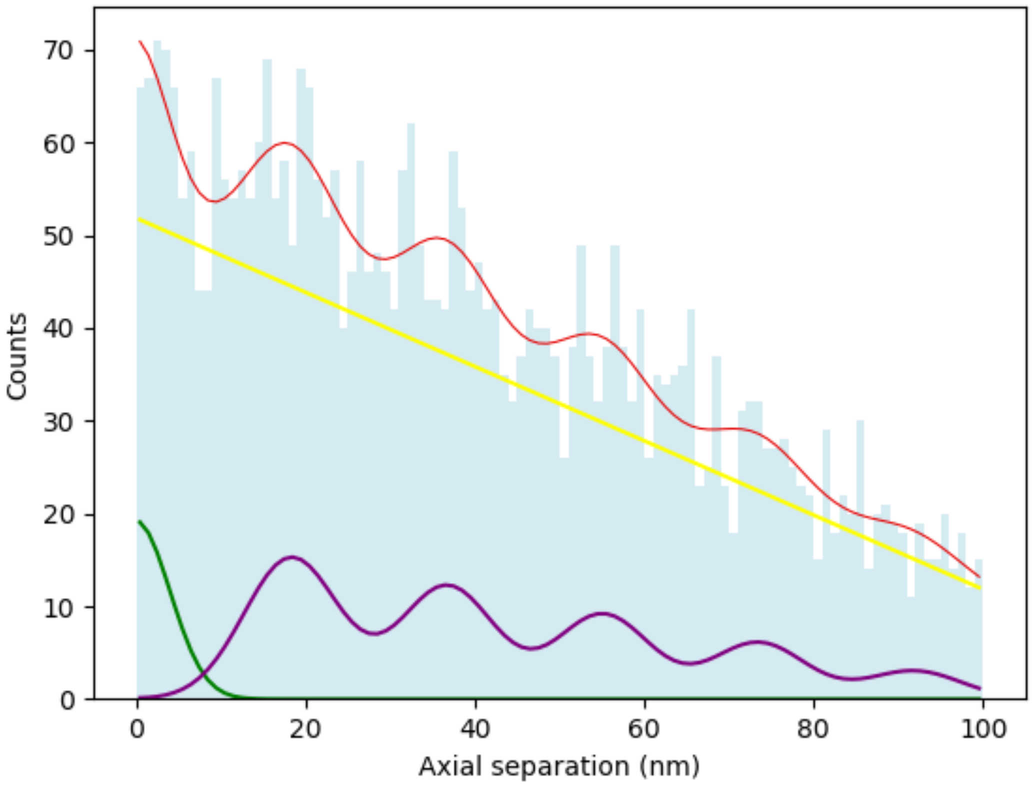


**Supplementary Fig. 5: Model components in the best-fitting model to the distribution of pairwise distances along the cell-axis between localisations of ACTN2 Affimer.** Red: optimised model (summed components). Yellow: linear term (background). Purple: peaks on linear repeat, height decreasing with distance. Green: localisation precision term (repeated localisations of one molecule).

**Supplementary Table 7: Akaike information and weightings (summing to one) of the relative likelihood of RPD models to be the closest to the true RPD, for pairwise distances along the cell-axis between localisations of the ACTN2 Affimer.** The models for a repeating distance along the cell-axis for ACTN2 are far more likely than the model for random distribution of localisations through the thickness of the Z-disk (linear fit), or the same but including repeated localisations (or clusters) of molecules. Furthermore, these fits all resulted in the same repeating distance for ACTN2 along the direction cell-axis, close to the 19.2 nm result measured previously by EM[7]. They may be grouped together to give a combined weighting of 0.97, compared with 0.03 for the random distribution model[4]. Combining the uncertainties for the linear repeat models gives a repeat of 18.4(1.0) nm.

| Axial RPD model | Fitted axial repeating distance | $AIC_c$ | Akaike weight |
|---|---|---|---|
| Linear repeat with 4 peaks (5-layer Z-disk) | 18.3(6) nm | 368.36 | 0.45 |
| Linear repeat with 5 peaks (6-layer Z-disk) | 18.4(5) nm | 368.97 | 0.33 |
| Linear repeat with 3 peaks (4-layer Z-disk) | 18.4(7) nm | 370.22 | 0.18 |
| Linear fit only (Random distribution) | N/A | 373.71 | 0.03 |
| Linear fit, with repeated localisations also allowed | N/A | 378.10 | 0.003 |

**Supplementary Table 8: Parameters and uncertainties for the RPD model with highest likelihood (linear repeat with 4 peaks, 5-layer Z-disk) for the distribution of pairwise distances along the cell-axis between localisations of the ACTN2 Affimer.**

| | | |
|---|---|---|
| **Linear term (background)** | Offset | 52(15) a.u. |
| | Slope | -3.8(1.6) a.u. $nm^{-1}$ |
| **Peaks on linear repeat, height decreasing with distance** | Linear repeat | 18.3(6) nm |
| | Amplitude of first peak | 202(228) a.u. |
| | Spread (broadening of peaks, s.d.) | 5.3(1.9) nm |
| **Localisation precision term** | Amplitude | 98(121) a.u. |
| | Spread (precision, s.d.) | 2.9(1.7) nm |

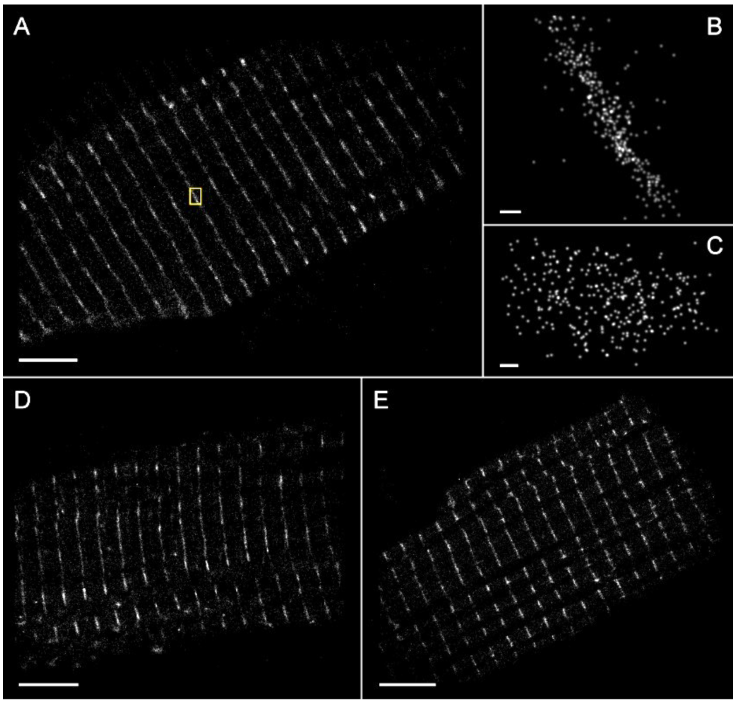


**Supplementary Fig. 6: 3D PALM on cardiac myocyte Z-disk proteins.** (A–C) mEos2:ACTN2, with cell axis illustrated (A). (B, C) Portion of a Z-disk (yellow box in A). (B) Higher magnification view of region in (A). (C) View onto the cell-transverse plane of the Z-disk portion (therefore viewed along the cell-axis). (D) mEos3.2:LASP2. (E) mEos3.2:MYPN. Scale bars: 5 µm (A, D, E), 200 nm (B, C).

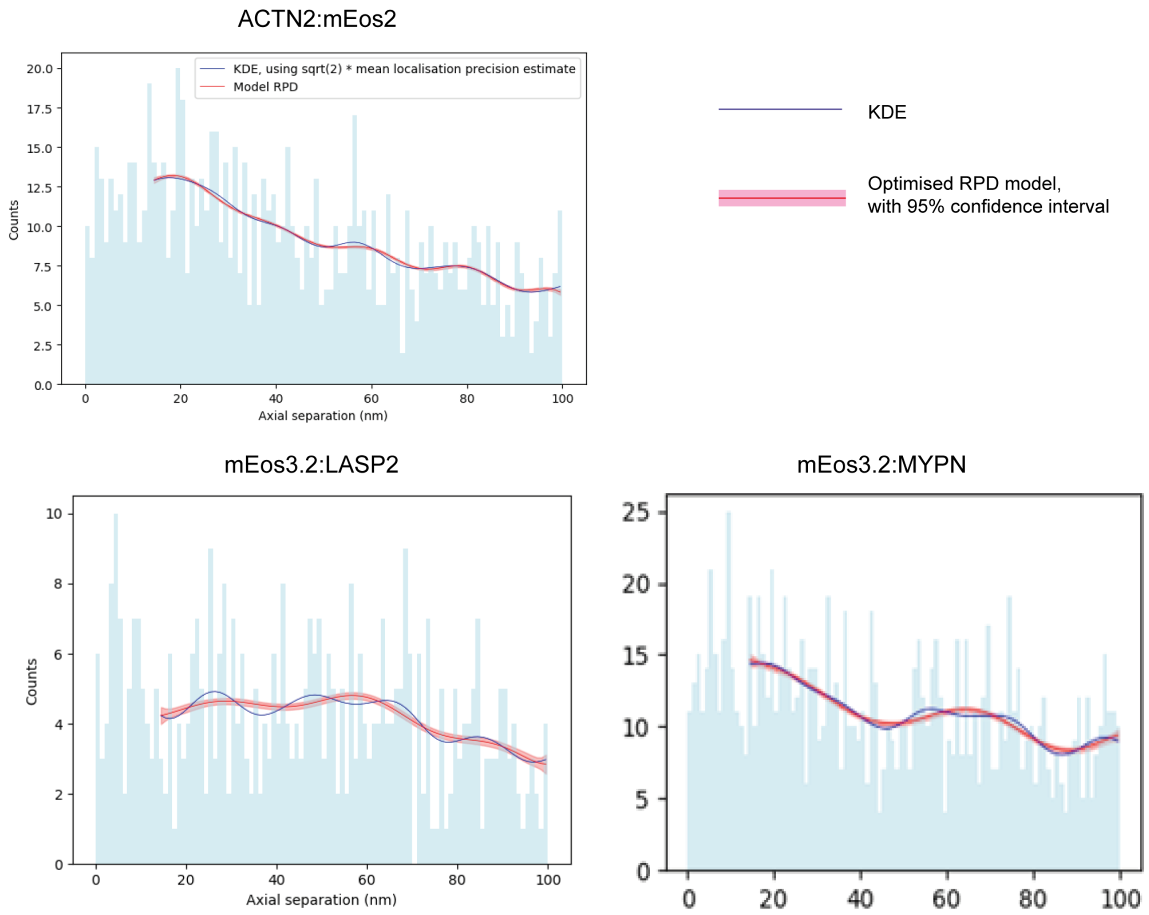


**Supplementary Fig. 7: Fits to axial separations in 3D PALM data on Z-disk proteins.** The distance histogram profiles are noisy as a result of the low fraction of proteins localised, and corresponding low numbers of relative positions. We obtained ~1400 ACTN2:mEos2 localisations per Z-disk within the localisation precision filter, or ~0.2% of likely instances of ACTN2. Therefore, we used a kernel density estimate (KDE) of the axial separations for model fitting, instead of the histogram bin counts. We used a Gaussian kernel with $\sigma$ = 4.8 nm ($\sqrt{2}$ × the mean estimated localisation precision). Distances below 3 × the kernel size were not fitted, to reduce the maximum effect on the estimate of the boundary at 0 nm to an error of 1%. Distances up to 100 nm + 3 × the kernel size were used to generate the KDE up to 100 nm, for a similar reason. We compared linear models, corresponding to random localisation distribution in the axial direction, and models containing a regular repeat for the position of the proteins, as done in the analysis of ACTN2 Affimer dSTORM data (**Fig. 2**, **Suppl. Table 7**). However, because of the smaller number of relative positions acquired, it was also necessary to allow the peak heights to vary independently, instead of fixing their ratios. Similarly to the ACTN2 Affimer dSTORM data, analysis of ACTN2:mEos2 localisations resulted in the correct identification of a regular repeating distance for ACTN2, this time at 20.20(8) nm. Models containing a regular repeat allowed the KDE of axial separations of mEos3.2:LASP2 and mEos3.2:MYPN to be fitted more closely than a linear (random positions) model, but resulted in a more heavily smoothened profile, with large uncertainties on parameter estimates (**Suppl. Table 9**). Therefore, PERPL analysis on this 3D PALM data has found very close to the correct repeating distance for ACTN2 (within 1 nm), but has not found a clear repeat for LASP2 and MYPN. However, the regularity in the peaks of the LASP2 KDE is striking, and more data and further analysis are required to investigate this more thoroughly.

**Supplementary Table 9: Parameters and uncertainties for the fitted RPD model (linear repeat with 5 peaks, 6-layer Z-disk) to the distribution of pairwise distances along the cell-axis between PALM localisations of Z-disk proteins.** All of these models were much more likely to be correct than a random distribution, according to AICc, but only the fit to the ACTN2 model resulted in an estimate of spread and parameter uncertainties that reasonably describe the regular density pattern considered.

**ACTN2:mEos2**

| | | |
|---|---|---|
| **Linear term (background)** | Offset | 11.4(5) a.u. |
| | Slope | -0.15(2) a.u. $nm^{-1}$ |
| **Peaks on linear repeat, height decreasing with distance** | Linear repeat | 20.20(8) nm |
| | Amplitude of peak 1 | 101(14) a.u. |
| | Amplitude of peak 2 | 90(18) a.u. |
| | Amplitude of peak 3 | 124(28) a.u. |
| | Amplitude of peak 4 | 162(36) a.u. |
| | Amplitude of peak 5 | 201(48) a.u. |
| | Spread (broadening of peaks, s.d.) | 8.8(2) nm |

**mEos3.2:LASP2**

| | | |
|---|---|---|
| **Linear term (background)** | Offset | 3.9(1.0) a.u. |
| | Slope | -0.14(51) a.u. $nm^{-1}$ |
| **Peaks on linear repeat, height decreasing with distance** | Linear repeat | 31(2) nm |
| | Amplitude of peak 1 | 151(5291) a.u. |
| | Amplitude of peak 2 | 283(1019) a.u. |
| | Amplitude of peak 3 | 359(1499) a.u. |
| | Amplitude of peak 4 | 419(1828) a.u. |
| | Amplitude of peak 5 | 10000(13038) a.u.* |
| | Spread (broadening of peaks, s.d.) | 15(6) nm |

* Since the $5^{th}$ peak at this repeating distance was some way from the distances fitted, its value is very uncertain and it reached the upper bound on the allowed parameter values during optimization.

**mEos3.2:MYPN**

| | | |
|---|---|---|
| **Linear term (background)** | Offset | 8(65) a.u. |
| | Slope | -0.38(46) a.u. $nm^{-1}$ |
| **Peaks on linear repeat, height decreasing with distance** | Linear repeat | 23(2) nm |
| | Amplitude of peak 1 | 468(2973) a.u. |
| | Amplitude of peak 2 | 277(635) a.u. |
| | Amplitude of peak 3 | 866(3821) a.u. |
| | Amplitude of peak 4 | 527(655) a.u. |
| | Amplitude of peak 5 | 1515(5909) a.u. |
| | Spread (broadening of peaks, s.d.) | 17(22) nm |

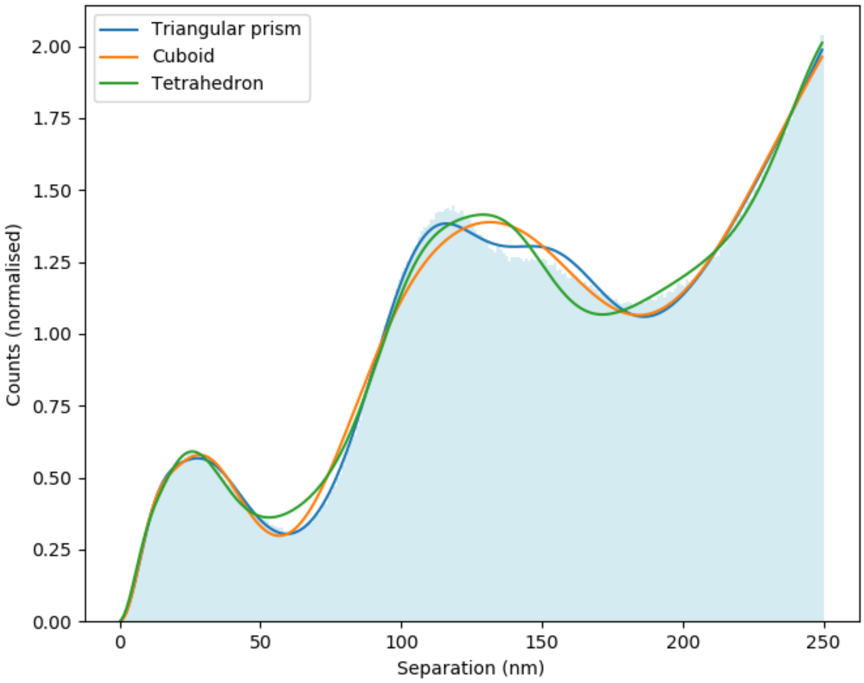


**Supplementary Fig. 8: RPD models fitted to the distance histogram of localisations of dye labels on a DNA origami structure.** The models included a square lattice pattern with broadening on the distances between lattice points, to match the apparent distribution on such a lattice of labelled structures in the reconstructed FOV. The models also included an isotropic 2D background model, since the data was more confined in Z than in XY. Two triangular prism models follow the same curve: in one the sides are all equal, in the other the length of the edges on the equilateral triangular faces was independent of the length of the connecting edges between them. In the cuboid model, the three edge lengths were independent of each other. In the tetrahedron model, the edge length on the equilateral triangular base was independent of the height of the pyramid.

**Supplementary Table 10: Akaike information and weightings (summing to one) of the relative likelihood of RPD models to be the closest to the true RPD, for the models in Suppl. Fig. 9.**

| RPD model | $AIC_c$ | Akaike weight |
|---|---|---|
| Triangular prism on square lattice (all sides equal)* | -1748.53 | 0.75‡ |
| Triangular prism on square lattice (unequal sides)* | -1746.33 | 0.25 |
| Cuboid on square lattice | -1477.75 | $1.2\times10^{-59}$ |
| Tetrahedron on square lattice | -1438.53 | $3.6\times10^{-68}$ |

* The side length in the model with all sides of equal length was 105.5(4) nm, while the two side-lengths in the unequal-sides model were 105(1) nm and 106(2) nm. These have therefore resulted in essentially the same model, but the AICc has penalised the model that allows unequal side-lengths for having an extra parameter, and results in the selection of the more parsimonious model with all sides equal.

‡ Selected model.

**Supplementary Table 11: Parameters and uncertainties for the most likely RPD model (triangular prism, equal sides) to the distribution of pairwise distances between DNA-PAINT localisations of labelled DNA-origami structures.**

| | | |
|---|---|---|
| **Localisation precision term** | Amplitude | 2.8(3) a.u. |
| | Spread (precision, s.d.) | 6.4(3) nm |
| **Inter-vertex distances** | Length of edges on triangular face | 105.5(4) nm |
| | Amplitude of contributions | 2.49(5) a.u. |
| | Spread (broadening of contributions, s.d.) | 19.9(3) nm |
| **Substructure at vertices** | Amplitude of contribution | 15.7(4) a.u. |
| | Spread (cluster size, s.d.) | 14.3(3) nm |
| **Square lattice arrangement of prisms*** | Square side length | 311(23) nm |
| | Amplitude of contribution | 325(125) a.u. $nm^{-1}$ |
| | Spread (s.d., due to the prism structure at each lattice point) | 72(11) |
| **Background term** | Gradient | $3.3(2)\times10^{-3}$ a.u. $nm^{-1}$ |

* These parameters are less well-estimated, since the relevant features reflecting the lattice spacing would be beyond the maximum fitted distance on the distance histogram. However, inclusion of the features in this way did allow the models to fit to the increase in the distance histogram from 200 to 250 nm.

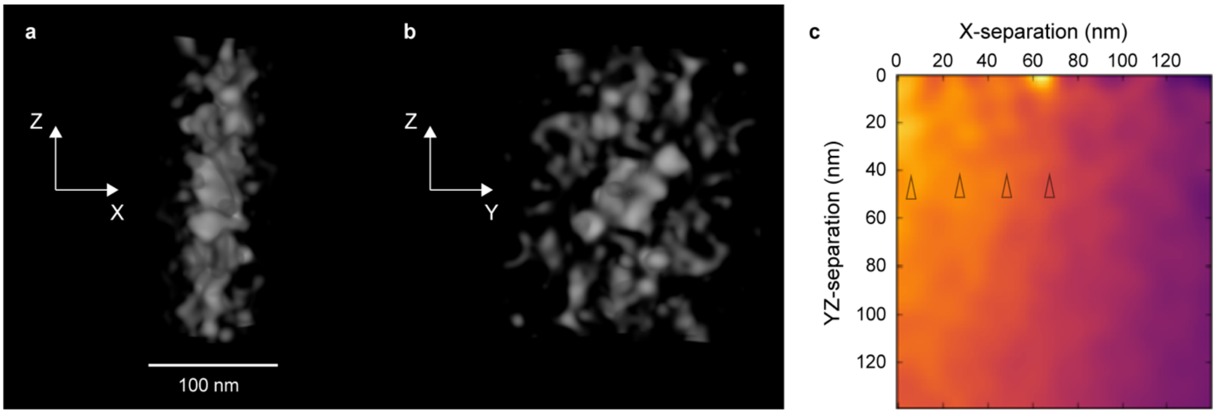


**Supplementary Fig. 9: 3D and 2D histograms obtainable in PERPL analysis.** 3D pairwise relative positions between cardiomyocyte ACTN2:mEos2 localisations plotted in a 3D histogram (**a**, **b**, with contrast stretching and Gaussian filtering; 3D projections in FIJI[8]), and in a 2D histogram (**c**, with Gaussian filtering). The axis of the cardiomyocyte points in the X-direction (see main text, **Fig. 2**); the confinement of ACTN2 to the thickness of the Z-disk is evident in the XZ view (a), while indications of the tetragonal lattice are seen in the YZ view (b). The ~20 nm repeat of ACTN in the axial direction also produces features in the 2D distance histogram (black triangles, c).

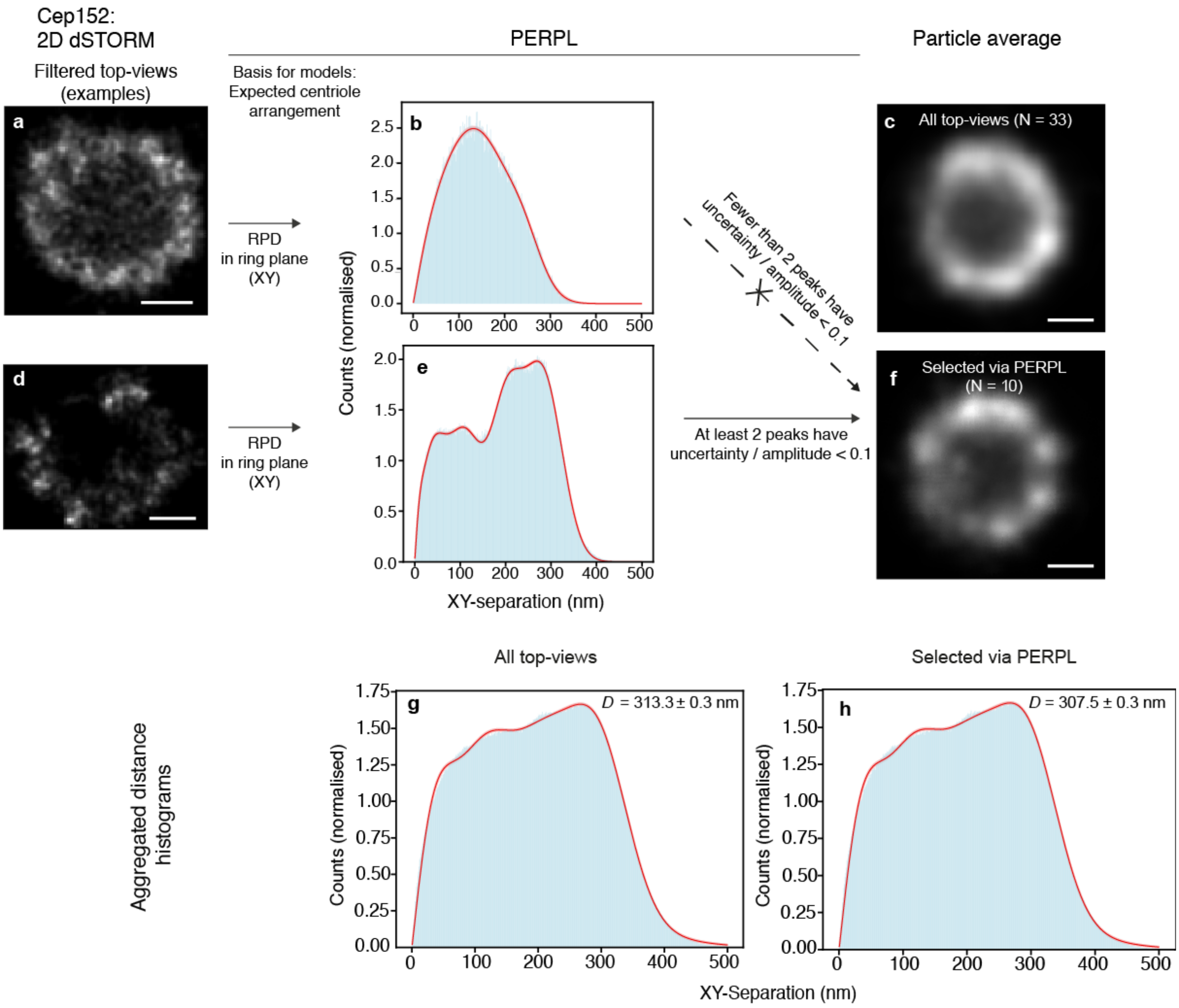


**Supplementary Fig. 20: Use of PERPL for particle selection.** (**a**, **d**) Examples of single dSTORM reconstructions of Cep152, filtered as top-views[9]. (**b**, **e**) Distance histograms and model RPDs (red), including 9-fold rotational symmetry, isotropic background, repeated localisations and cluster size (see **Fig. 1**). 95% confidence intervals (pink, **b**, **e**, **g**, **h**) are narrow and not clearly visible at this scale. (**c**) Result of averaging all 33 filtered Cep152 images. (**f**) Result of averaging the 10 Cep152 images remaining, after analysing and thresholding for uncertainty (s.d.) of peak amplitudes in the nine-fold symmetry component of the model fit. There are four peaks representing inter-vertex distances in the nine-fold symmetry component of the model, similar to the model illustrated in **Fig. 1**. The background term was not used. Because analysis was per particle, and particles had missing vertices, we allowed the amplitudes of these four peaks to be four separate variables. We selected only particles where the uncertainty (1 s.d.) of the estimated peak amplitude was < 0.1 × peak amplitude for at least two of the peaks, and obtained an improved particle average, showing distinct clusters and a rounder structure for the complex. (**g**, **h**) Distance histograms aggregated over all or selected particles and model RPDs, with diameter (*D*) estimated from the least-squares fits. PERPL analysis also results in diameter measurements without requiring successful particle averaging. The diameters estimated from all particles and from the selected particles were within 3 nm. Use of the model of **Fig. 1** with constrained ratios of inter-vertex distance did not result in good fits here, which may be a result of missing distances in the small number of particles used, imperfection in the orientation of 'top-view' centriole images, or deviation of cluster shape from the Gaussian assumption.

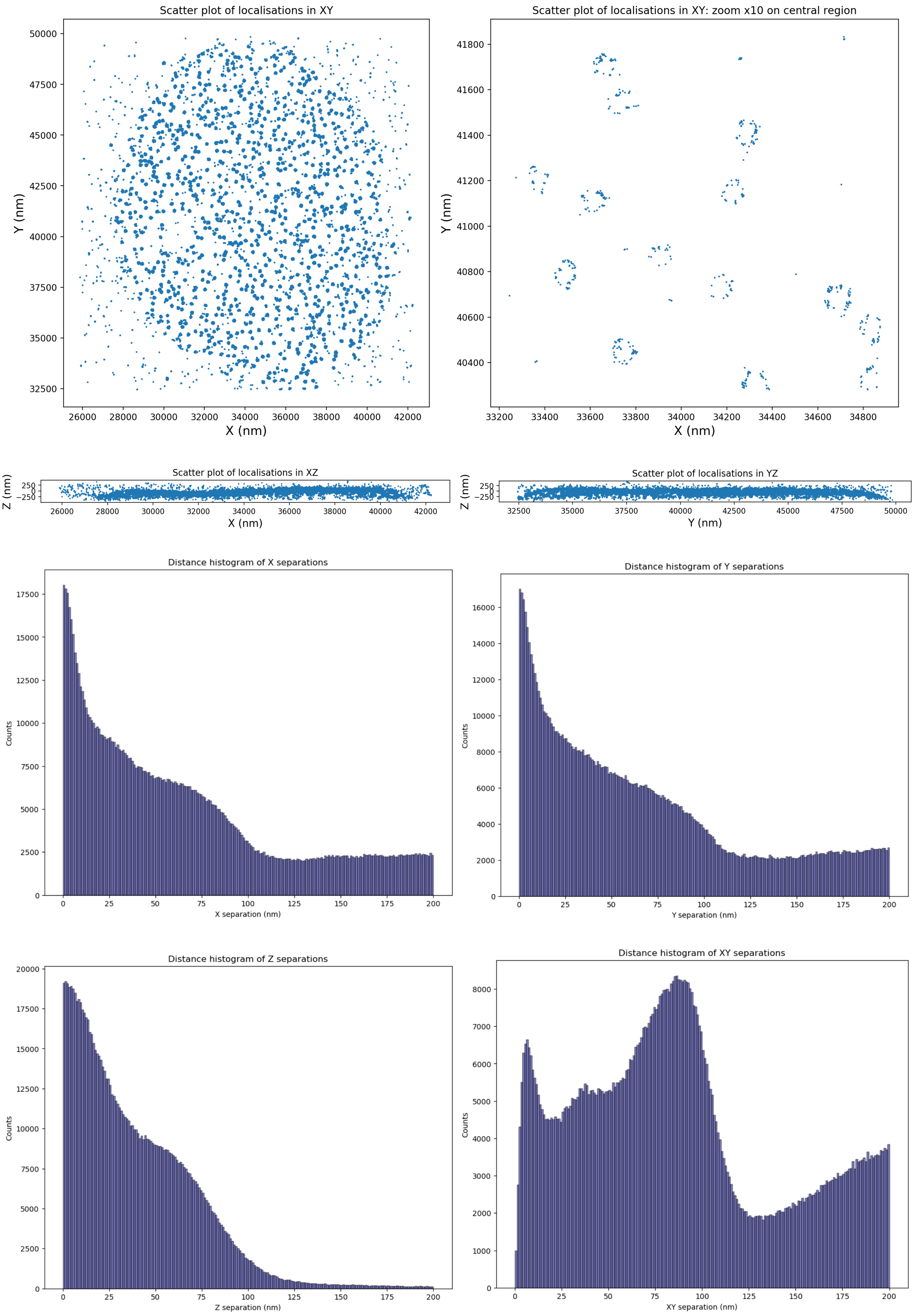

Scatter plot of localisations in XY
Y (nm)
X (nm)
Scatter plot of localisations in XY: zoom x10 on central region
Y (nm)
X (nm)
Scatter plot of localisations in XZ
Z (nm)
X (nm)
Scatter plot of localisations in YZ
Z (nm)
Y (nm)
Distance histogram of X separations
Counts
X separation (nm)
Distance histogram of Y separations
Counts
Y separation (nm)
Distance histogram of Z separations
Counts
Z separation (nm)
Distance histogram of XY separations
Counts
XY separation (nm)

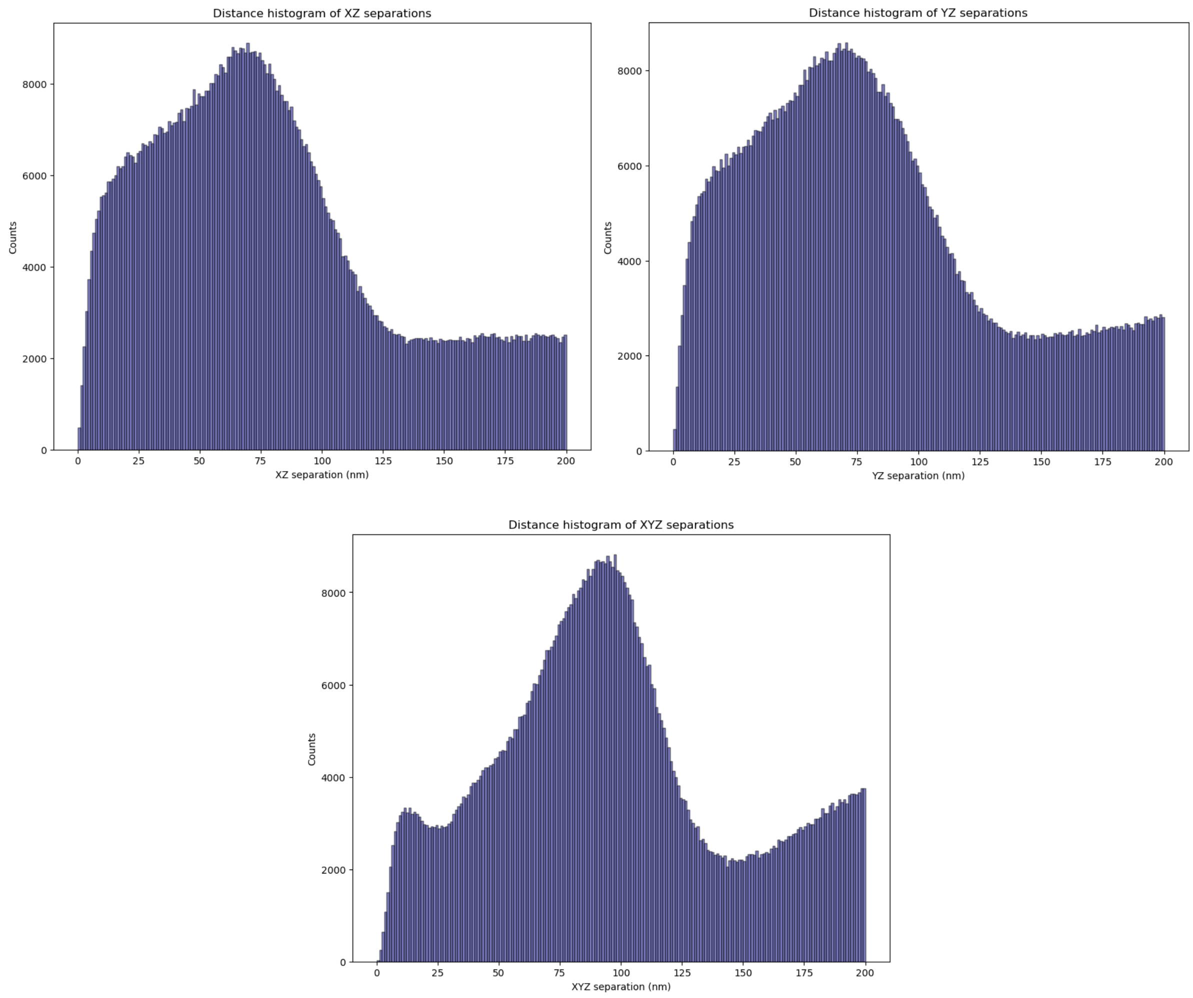


**Supplementary Fig. 11: Scatterplots of the localisations and distance histograms in various directions saved when *relative_positions.py* is run on a list of localisations.** These are placed into an html report for the user, as well as being saved individually.

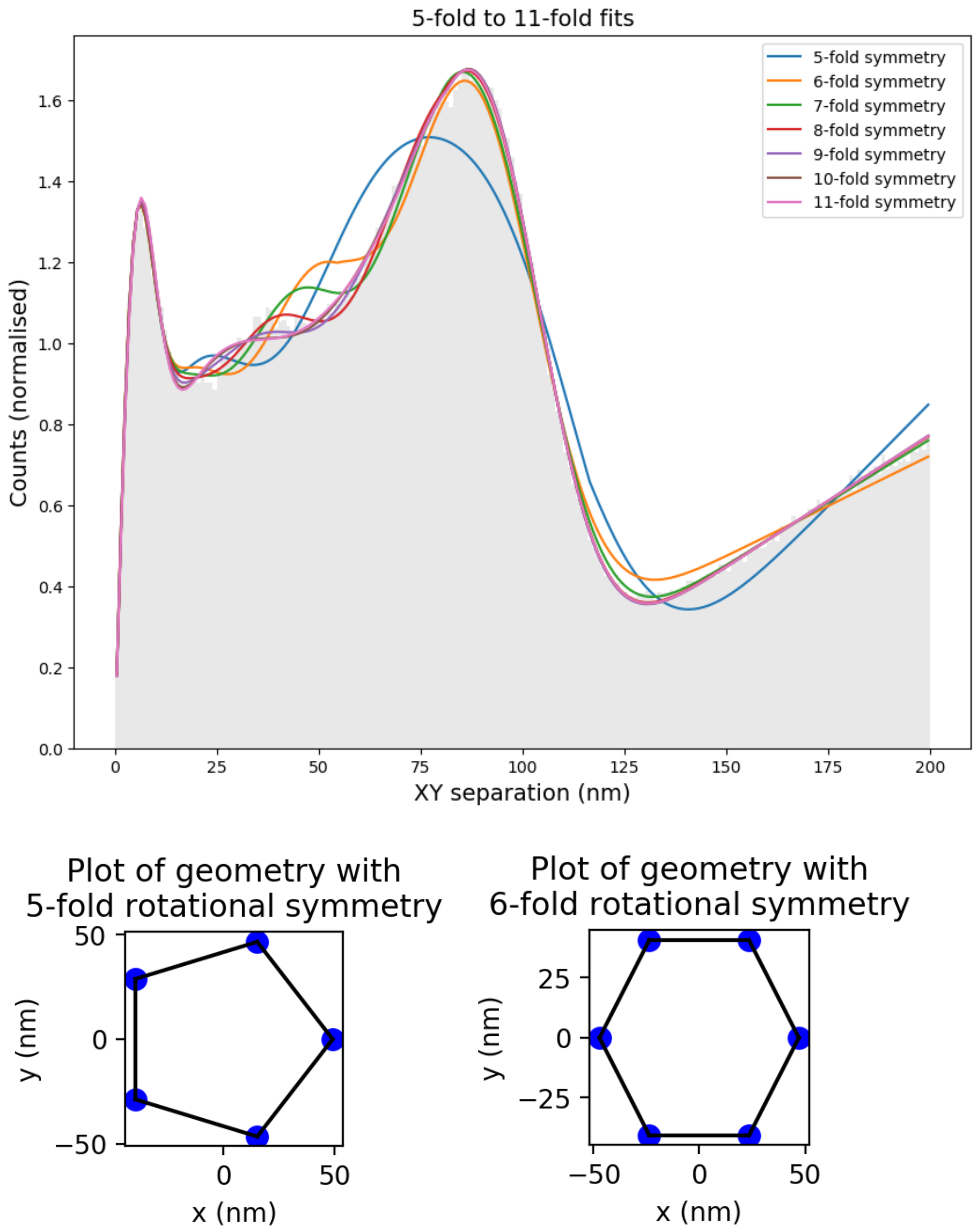


**Supplementary Figure 12: Examples of plots saved and contained in the html report when *rot_2d_symm_fit.py* in Supplementary Software is run on output data from *relative_positions.py*.** The optimized models for each order of symmetry are plotted on the XY-distance histogram and simple plots of the optimised geometry at each order of symmetry are generated. The html report from *rot_2d_symm_fit.py* also contains the AICc values, relative likelihoods, and parameter estimates and uncertainties for each modelled order of symmetry. Additionally, it contains descriptions of the model parameters and their initial guesses and bounds as used by the least squares fitting function.
